# On the impact of re-mating and residual fertility on the Sterile Insect Technique efficacy: case study with the medfly, *Ceratitis capitata*

**DOI:** 10.1101/2023.08.17.552275

**Authors:** Yves Dumont, Clelia Felice Oliva

## Abstract

The sterile insect technique (SIT) can be an efficient solution for reducing or eliminating certain insect pest populations. It is widely used in agriculture against fruit flies, including the Mediterranean fruit fly (medfly), *Ceratitis capitata*. The re-mating tendency of medfly females and the fact that the released sterile males may have some residual fertility could be a challenge for the successful implementation of the SIT. Obtaining the right balance between sterility level and sterile male quality (competitiveness, longevity, etc) is the key to a cost-efficient program. Since field experimental approaches can be impacted by many environmental variables, it is difficult to get a clear understanding on how specific parameters, alone or in combination, may affect the SIT efficiency. The use of models not only helps to gather knowledge, but it allows the simulation of a wide range of scenarios and can be easily adapted to local populations and sterile male production.

In this study, we consider single- and double-mated females. We first show that SIT can be successful only if the residual fertility is less than a threshold value that depends on the basic offspring number of the targeted pest population, the re-mating rates, and the parameters of double-mated females. Then, we show how the sterile males release rate is affected by the parameters of double-mated females and the male residual fertility. Different scenarios are explored with continuous and periodic sterile males releases, with and without ginger aromatherapy, which is known to enhance sterile male competitiveness, and also taking into account some biological parameters related to females that have been mated twice, either first by a wild (sterile) male and then a sterile (wild) male, or by wild males only. Parameter values were chosen for peach as host fruit to reflect what could be expected in the Corsican context, where SIT against the medfly is under consideration.

The obtained results confirmed that the use of ginger aromatherapy is an essential step for SIT success against medfly. Our work also highlights the importance of estimating the duration of the refractory period between matings and how it may change when a wild (or sterile) female mates first with a sterile (or wild) male. Further, we show the importance of parameters, like the (hatched) eggs deposit rate and the death-rate related to all fertile double-mated females. In general, re-mating is considered to be detrimental to SIT programs. However, our results show that, depending on the parameter values of double-mated females, re-mating may also be beneficial for SIT. Our model, being generic, can be easily adapted to different contexts and species, for a broader understanding of release strategies and management options.

## 1 Introduction

The Sterile Insect Technique (SIT) is an established autocidal control method used effectively against several agricultural pests in many countries worldwide. Some of the major operational programs include the Mediterranean fruit fly (medfly) *Ceratis capitata*. SIT relies on a continuous mass production of the targeted insect, the sterilization of males (or both males and females, according to the species) as pupae or adults, using ionizing radiation, and their repeated and massive releases in the field that result in a progressive decay of the targeted pest population [1, 2].

For fruit flies, ionizing radiation is mostly used to sterilize males as part of the operational programs. However, in order to sterilize only males, and also from an economic point of view for large-scale releases, it is important, to separate males and females at an early stage. To this end, a genetic sexing strain (GSS) medfly [3], based on the temperature sensitivity of females, has been developed in the nineties. Since then, it has been improved and used in many places around the World [3]. Unfortunately, for massive production, maintaining the GSS-strain is quite technical, such that it cannot always be used in all SIT programs. However, improvements are under study, see for instance [4]. SIT is also never used alone, but it is the primary control program in several operational integrated pest management (IPM) programmes that may use additional suppression methods, like insecticides and mass-trapping, or other cultural practices, etc. See for instance [5].

An alternative approach of SIT, a genetically engineered sterile strain of medfly (*C. capitata*), has been developed and tested for population reduction on a small scale [6]. Other suppression approaches include the incompatible insect technique (IIT), which uses the bacterial endosymbiont Wolbachia [7], has been considered against Medflies [8]. In this study, we considered only the approach based on sterilization with irradiation.

Conceptually, the Sterile Insect Technique is simple, but its efficient deployment requires understanding and tailoring several technical, logistical, ecological, and biological parameters, as well as socioeconomic elements [1]. Although the use of SIT against medflies is widespread (it is used in South and Central America, USA, Australia, South Africa, etc.), in Europe it is only operational in Spain and Croatia. France is developing an SIT research pilot project to determine the optimal conditions for the deployment of the SIT on Corsica Island to protect stone fruits and clementine production (CeraTIS Corse project).

The Mediterranean fruit fly is the dominant fruit fly pest on citrus and deciduous fruit in Corsica, where currently, todays treatments still rely mostly on the use of pesticides. The growing demand for more environmentally friendly approaches, together with the potential future unavailability of chemical substances, triggered a pilot research project to integrate SIT in the control strategies. Most Corsica orchards have challenging initial ecological conditions (high density of flies, crop areas surrounded by rural settings, and wild host plants for medflies). Designing an efficient and cost-effective suppression program requires a good understanding of the biological and technical parameters that impact its success. As part of this effort, modeling the effect of sterile males’ residual fertility, female re-mating and release frequency on release ratios is essential.

SIT impacts the reproduction of the targeted insect. Hence, its success depends on the ability of the released sterile males to find a lek, to perform courtship, to be selected by wild females, to successfully copulate and inseminate females with their sterile sperm, while eliciting effective female refractoriness to further re-mating by wild males. Medflies are considered to have a complex courtship behavior that could make SIT less efficient [9]. Nevertheless, the SIT is a well established method of suppressing the population of medflies and has been proven to be highly efficient and economically viable in many applications [10]. Modeling, model analysis, and simulations can be helpful in highlighting the positive or negative impact of specific parameters and assist in formulating an optimal release strategy in the field.

SIT has been modeled since the early work by Knipling [2]. Various discrete and continuous models have been developed of varied complexity depending on the number of stages in the life of the targeted pest/vector: for instance discrete models [11, 12, 13], semi-discrete models [14, 15, 16] or continuous models [17, 18, 19], using sometimes tools from the control theory [20, 21, 22] to optimize the release protocols. Many SIT models consider sterile males to be fully sterilized, although this is rarely achieved. Indeed, reaching full sterility requires a very high dose of radiation [23], which often impairs the quality of males to a level that is not acceptable. However, if males are not fully sterile, meaning that a fraction of their sperm is still fertile. As a result, a small fraction of the sterile males will inseminate wild females that will produce viable eggs. This fraction is called residual fertility. One could argue that releasing millions of almost-sterile males is equivalent to releasing a certain percentage of fertile males.

Residual fertility is easy to assess and is part of the quality control of every SIT program (interested readers are referred to the manual published by the International Atomic Energy Agency (IAEA) [24] for medfly SIT procedures). For *C. capitata*, a dose of 140Gy is required to achieve full sterility, as in the medfly management program in Argentina [25]; however, this dosage can also negatively affect male performance and therefore be detrimental for SIT operations [26, 27]. On the other hand, in an ongoing operational SIT program against medflies in the region of Valencia, Spain, the induced average sterility reached 98.87 ± 0.55% with an irradiation dose of 100 Gy, leading to a residual fertility of 1.13 ± 0.55% [5]. This program is still running with good results.

Therefore, it is important to know the threshold above which residual fertility can have a negative impact on SIT program performance and how to estimate it. In a recent study, on a different fruit fly species [16] and considering single mating only, we investigated this question using a very simple and minimalistic model.

The result was straightforward: the percentage of residual fertility (RF), *ε*, must be lower than 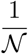, where 𝒩 is the basic offspring number, also called the basic reproduction number, which is related to the reproductive potential of the pest population. Clearly, for a wild population with a large 𝒩, the constraint on RF can be strong, thereby significantly reducing its acceptable level for successful SIT. A similar result was obtained by Van den Driessche [11] using discrete models. In this study, we explore a more complex model, with single and double mating, to verify these results, with a particular focus on medfly. Furthermore, we considered the important parameter of re-mating and whether there is a different response when wild females mate with sterile males as opposed to wild males.

Female medflies are facultatively polyandrous [28, 29]. After mating, medfly females typically exhibit an average refractory period of two and a half days [30]. Their propensity to remate can be triggered by the courtship behavior of the male, sexual performance, and amount of ejaculate transferred [31, 32, 33]. The likelihood of re-mating might (as evaluated under laboratory conditions) increase in situations of high availability of oviposition substrate or highly male-biased sex ratios, which is the case of male-only release SIT programs [30, 34]. Medfly females possess 2 spermathecae, spherical organs that store sperm after insemination, and a fertilization chamber that serves as a functional third spermatheca [35]. Having more than one spermatheca could be seen as advantageous in case of multiple insemination. In most insects, the sperm from different males is mixed within the spermathecae, therefore not physically allowing for preferential selection during egg fertilization. However, the fertilization chamber in medflies allows a second male to remove sperm from the previous one [35]. Sperm precedence of the second male mating a female tends to win in fertilizing eggs (second male contribution greater than 0.5) [28, 36, 37, 33, 38]. In [39], the authors studied double-mated females and show how the sperms of the first and the second mating are stored, meaning that both are able to be used to fertilize the eggs. In [38, 40], the authors show that re-mating increases the fitness of females, while Katiyar et al. [37] showed that copulation order between fertile and sterile males impacts the fitness of females, with a precedence of the second sperm. Lee et al. [36] reported that male genotype, copulation order and genotypic differences may affect the variation in sperm precedence. From the SIT point of view, this not necessarily a good news. In the rest of the paper, for sake of simplicity, we will consider single- and double-mating, but our reasoning could be extended to more than two matings. In the present study, we investigated the mentioned two parameters, namely residual fertility and remating, can impact the required release ratios under different situations. Most medfly SIT programs involve biweekly releases, some with daily releases. Although release frequency has a direct impact on the program efficiency, it is also useful to understand how this impact is affected by biological parameters. More precisely, as most programs have implemented the addition of ginger root oil (GRO) aromatherapy to enhance sterile male attractiveness [30], we also included a comparison of competitiveness parameters for treated or untreated males. In the rest of the paper, we will mostly consider data related to the V8 (Vienna-8) strain obtained from laboratory colonies maintained by the Seibersdorf laboratory of IAEA (International Atomic Energy Agency, Vienna, Austria). The V8-strain is a GSS-strain: it allows to separate easily males and females. In [41], the authors showed that 99% of V8 strain copulations resulted in sperm transfer, and that the V8-strain has a good re-mating potential.

The paper is organized as follows: in Section 2, we build the continuous SIT model with re-mating and residual fertility. We also derive some theoretical results before extending the model to periodic impulsive releases. In Section 3, we provide numerical simulations related to continuous and periodic releases, highlighting different re-mating cases. We discuss the results in section 4. Finally, in Section 5, we derive some conclusions and perspectives.

## 2 Material and methods

Assuming a large population of medflies and the fact that all interactions between individuals and all biological processes occur simultaneously, we consider a continuous modeling approach, using a system of ordinary differential equations to model our biological system. In the forthcoming model, we consider several compartments (see the compartmental diagram in Fig. 1, page 6): flies in immature non-flying stages, *A*, encompassing larvae and pupae; wild males, *M* ; fertile wild females, *F*_*W*_ ; sterile females, *F*_*S*_; fertile wild females re-mated with a sterile male, *F*_*W S*_; sterile females re-mated with a wild male, *F*_*SW*_ ; fertile wild females re-mated with a wild male, *F*_*W W*_ ; double-mated sterile females, *F*_*SS*_; “almost” sterile males, *M*_*S*_. We also consider that our biological system is isolated, such that there is no migration of fruit flies. The females *F*_*S*_ and *F*_*SS*_ are assumed to be fully sterile.

**Figure 1:**
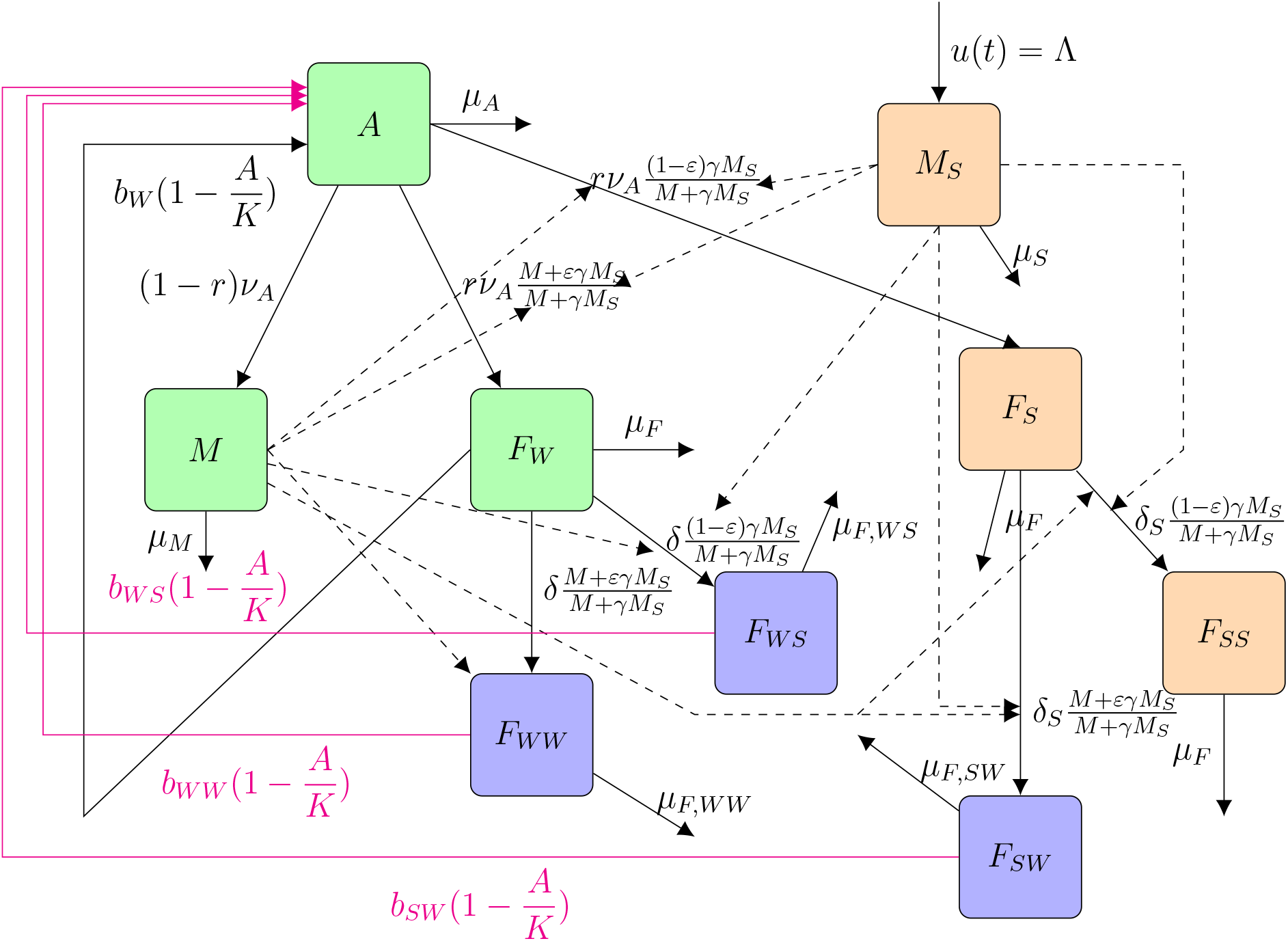
Flow diagram of model (1): the green squares represent the (fully) wild population; the blue squares represent the double-mated fertile females; the orange squares represent the sterile population.

We consider different birth-rates and also death-rates for each (fertile) female compartments, that is *b*_*W*_, *b*_*W W*_, *b*_*W S*_, *b*_*SW*_. In particular, following [40], we have *b*_*W W*_ *≥ b*_*W*_ and *μ*_*F,W W*_ < *μ*_*F*_ : re-mated females have greater longevity and higher productivity than one-time maters. For the parameters *b*_*_, related to *F*_*SW*_ and *F*_*W S*_, we have some scarce knowledge[37, 36], but nothing on *μ*_*_. In the numerical part, we will make several simulations to show the importance of knowing these parameters.

We model the residual fertility like in [16, 42, 43]: we assume that in the sterile males population there is a small proportion, *ε*, of fertile sperms, such that at the population-level, we assume that a proportion of *ε* sterile males is able to fertilize females. All variables and parameters are described in Table 1, page 5. As seen in diagram 1, the birth rates are impacted by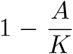, where *K* is the carrying capacity (i.e. maximum number of larvae/pupae for all fruits) of the host(s). Since the average mating rate at first mating was shown to be similar for wild and V8 sterile males (0.67 and 0.64, respectively, [41]), we consider equal chance for wild and sterile males to mate. Without residual fertility, the impact of SIT is modeled by the term 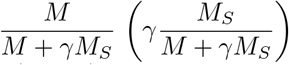 which represents the probability for a sexually mature female to mate with a wild (sterile) male and enter one of the mated female compartments, *F*_*W*_ (*F*_*SW*_, *F*_*W*_) or *F*_*W W*_ (*F*_*S*_). Male residual sterility is modeled by considering that a proportion, *εM*_*S*_, of sterile males is fertile, such that emerging immature females will become fertile with a probability of 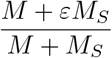 or they will become sterile with a probability of 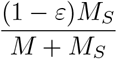.

**Table 1:**
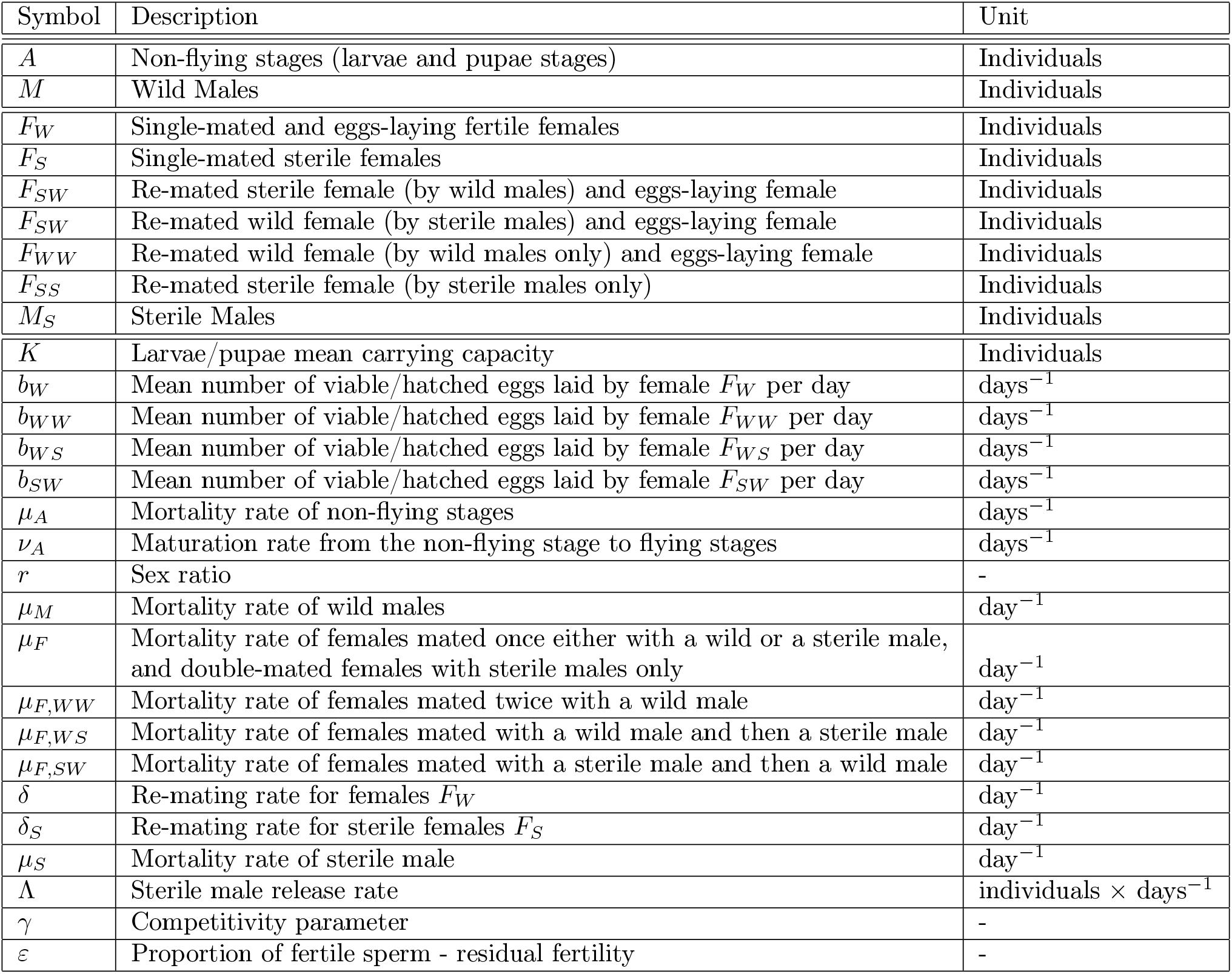
Description of parameters and state variables of model (1)

All linear terms represent either transfer rates, like *ν*_*A*_ and *ν*_*Y*_, from one compartment to another, or death rates, *μ*_*A*_, *μ*_*M*_, *μ*_*F*_ and *μ*_*S*_: see Table 1, page 5. We now explain further the re-mating component that is also specifically investigated in this study.

Host fruit affect the rate and duration of development, in this study we considered values from stone fruits when available, as peaches, nectarines, and apricots are the most common fruits in Corsica, together with clementines. Peaches [44] and nectarines [45] appear to be very suitable fruits for the development of *C. capitata*, while clementine is less favorable for immature development [44] (see table (1) and (2) for the parameters used and their value). Developmental data also vary with season (temperature and fruit phenology), but those variables are not included in our model.

We take into consideration male and female’s multiple mating capacity (re-mating). This parameter has been repeatedly studied in the laboratory with varying estimations. However, analysis of field-sampled females progeny showed less than 28% [46] or 50% [28] multiple mating. It is then possible to roughly estimate a percentage of female daily re-mating proportion: in average, between 7% and 12%.

This is represented, in our model, by the parameters *δ* and *δ*_*S*_, where where 1*/δ* and 1*/δ*_*S*_ represents respectively the average refractory periods for the wild female, *F*_*W*_, and the sterile female, *F*_*S*_. In general, some studies (for example [47]) showed that wild females that have mated with sterile males, *F*_*S*_, have a tendency to re-mate more often, such that we will consider *δ*_*S*_ *≥ δ ≥* 0. When a female *F*_*W*_ re-mates, then she re-mates with either a wild male or a sterile male to enter either the compartment *F*_*W W*_ at the rate 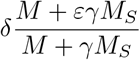 or the compartment *F*_*W S*_ at the rate 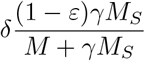. This is similar for a sterile female that can re-mate with a wild male and thus enter in the compartment *F*_*SW*_ at the rate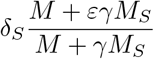.

The addition of ginger root oil (GRO) to sterile adult males is now a common process that increases the sterile males competitiveness. It has also been showed to reduce re-mating tendency, leading to similar percentages of re-mating whether females are mated first to wild or sterile males [48]. In this study we analyse the effect of GRO treatment on the release ratios.

The released number of sterile males may vary in time. Thus, we consider a release rate *u*(*t*) *≥* 0. However, for sake of simplicity and in order to go as far as possible in the qualitative analysis, we will mainly consider the constant and continuous release case, *u*(*t*) *≡* Λ_*S*_. However, it is also possible to consider periodic impulsive releases, i.e.

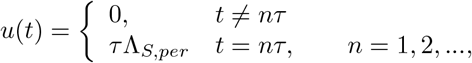

where *τ* is the given days between two consecutive releases, like 3 or 7 days.

The compartmental diagram in Fig. 1, page 6, can be translated into the following mathematical model

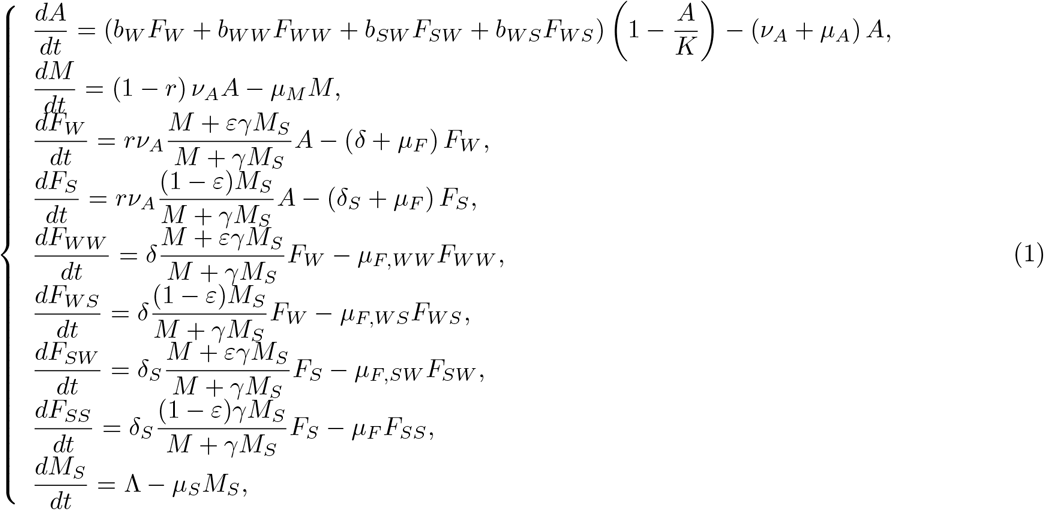

with non-negative initial conditions. Assuming *t* large enough, we may assume that *M*_*S*_ has reached its equilibrium,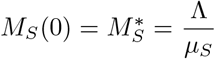. In addition, the *F* _*SS*_ -equation (1) 8 being not involved in the dynamics of the double-mating model (1), studying system (1) is equivalent to study the following system

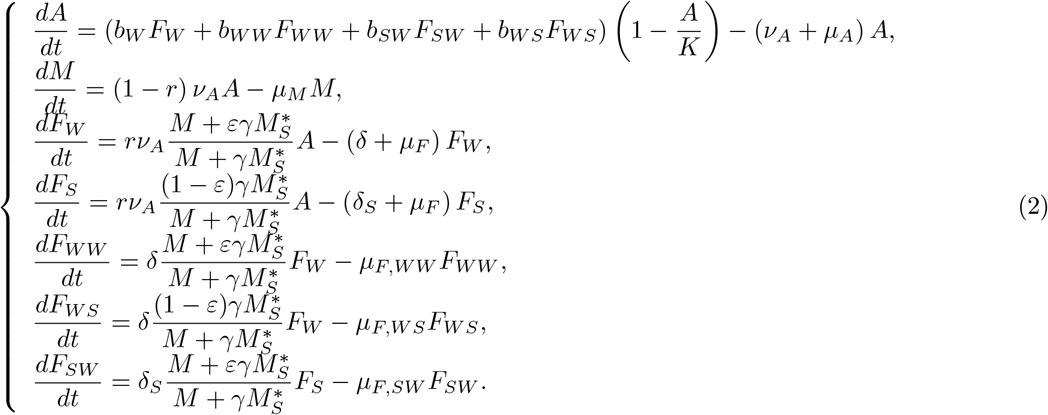

Even if system (2) is not cooperative, it can be studied using the monotone theory approach [49]. Indeed, since 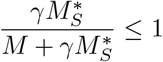 in equations (2)_4,6_, it is straightforward to show that system (2) is upper bounded by the following monotone cooperative auxiliary system [49]:

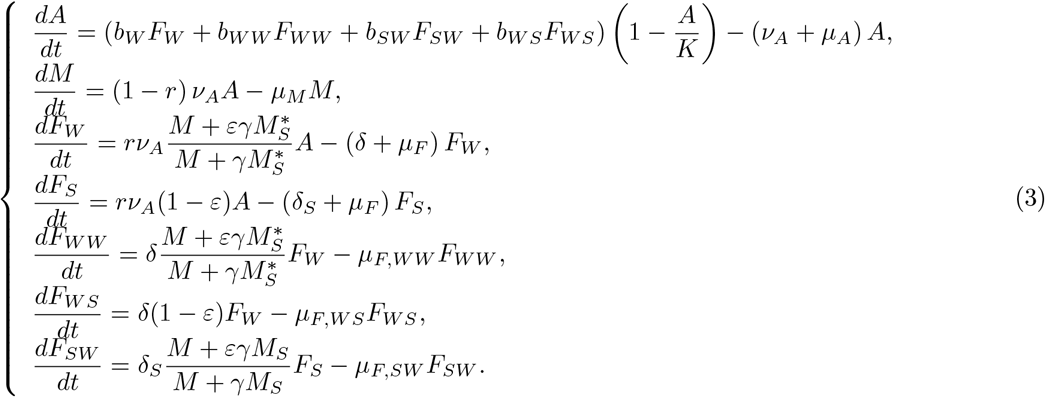

We will see later how useful is auxiliary system (3).

The following lemmas show that system (3) is mathematically and biologically well posed: it remains positive and bounded.

### Lemma 1

*Let M*_*S*_ *be a non-negative, piecewise continuous function on* R_+_ *and assume non-negative initial data. The solution to the Cauchy problem associated with* (3) *exists on, is unique, continuous and* 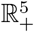 *piecewise continuously differentiable. This solution is also forward-bounded and remains non-negative. It is positive for all positive times if F* (0), *Y* (0) *or F* (0) *is positive*.

It is also straightforward to show that the set 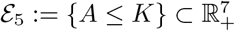 is forward invariant for (2), and any trajectory enters it in finite time.

### 2.1 Qualitative analysis without SIT releases

Without sterile male releases, we recover from any of (1), (2) or (3) the model of the dynamics of the natural/wild pest/vector population

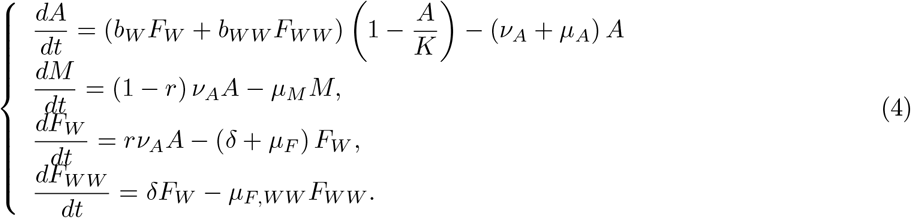

We briefly study model (4). We define the basic offspring number related to model (4) as follows

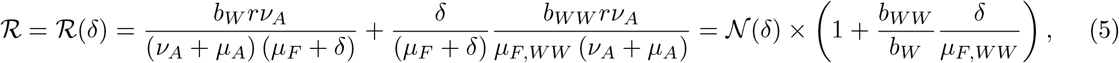

where

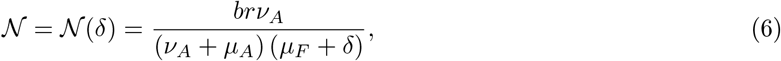

#### Remark 1

*The parameter ℛ represents the average number of (female) offspring a single-mated female and a double-mated female can produce during their life time. The parameter 𝒩 represents the average number of (female) offspring a single-mated female can produce during her life time*.

Through straightforward calculations and using the theory of monotone cooperative system [49], we show the following result:

#### Theorem 1 ([50])

*System* (4) *defines a forward dynamical system in* 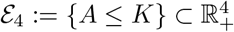. *In addition*

- *if ℛ* < 1, *then* 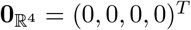 *is globally asymptotically stable on E*_4_.
- *if ℛ* > 1, *then a positive equilibrium* **E** *exists where*

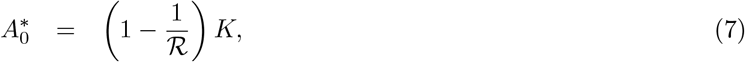

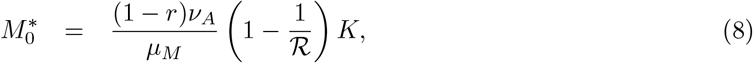

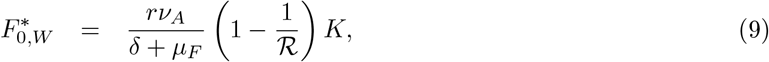

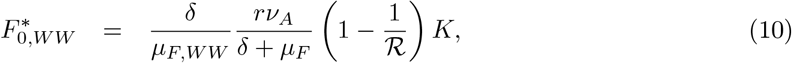

*Furthermore*, **E** *is globally asymptotically stable on E*_4_ *\ {*(0, *M*, 0, 0) : *M ≥* 0*}*.

**Proof**. See Appendix A, page 30.

#### Remark 2

*It is interesting to notice that 𝒩 ≤ ℛ, so that 𝒩* > 1 *implies ℛ* > 1, *and ℛ* < 1 *implies 𝒩* < 1.

### 2.2 Some theoretical results on the SIT system

For the rest of the paper, we assume that ℛ> 1. SIT induces a strong Allee effect [51] by reducing mate finding probabilities such that a population level extinction threshold exists and such that we get bi-stability between 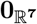 and a positive equilibrium, 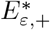, when sterile males releases are not too large. This effect is particularly useful when the pest population is either (very) small or invading the domain, or to consider a long term SIT strategy using first massive releases, and then small releases, as explained in [51, 16]. Indeed, massive releases will drive the pest population into a subset that belongs to the basin of attraction of 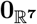, related to a given size for the small releases, such that we can switch from massive to small releases to keep the pest population as small as needed and also to drive it slowly, but surely, towards elimination, i.e. 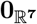. In contrast, once SIT is used, it has to be maintained: if, for any reason, SIT is stopped, then the Allee effect is lost and the wild population will rise again.

In the following Lemma, we derive a condition on residual fertility which ensures that equilibrium 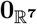 is Locally Asymptotically Stable, i.e. that the strong Allee effect exists.

#### Lemma 2

*When ε ≤ ε*_max_, *where ε*_max_ =

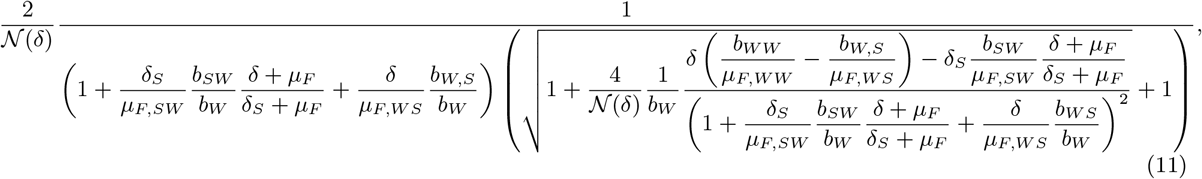

*then* 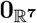 *is always Locally Asymptotically Stable (LAS) for system* (2). *It is unstable, otherwise*.

**Proof**. See Appendix B, page 32.

From the previous Lemma, we deduce that if RF is too large, i.e. *ε* > *ε*_max_, there is no strong Allee effect, whatever the size of sterile male releases.

The upper bound for the residual fertility, *ε*_max_, given in (11), is particularly interesting, because it does not only depend on re-mating parameters *δ, δ*_*S*_, but also on the (hatching) eggs deposit rates and the death rates for each type of Females, *F*_*W*_, *F*_*W S*_, *F*_*SW*_, and *F*_*W,W*_.

#### Remark 3

*Without remating, i*.*e. δ*_*S*_ = *δ* = 0, *we recover the condition ε ℛ* (0) *≤* 1, *as obtained in [16], for instance*.

In formula (11), the term

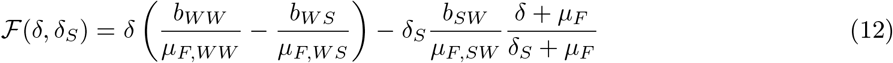

is particularly interesting because, depending on the previous parameters, it can be either negative, or nonnegative. Thus, when *ℱ* < 0, then re-mating is reducing the impact of SIT, while when *ℱ≥* 0, re-mating is neutral or beneficial for SIT. For instance, if *δ* = *δ*_*S*_ = 0, then *ℱ* = 0. If *δ*_*S*_ = *δ* (equal re-mating), then

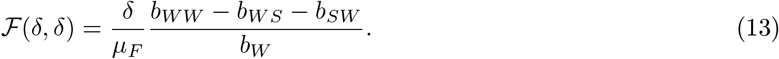

Hence, if *b*_*W W*_ = *b*_*W S*_ + *b*_*SW*_ then ℱ = 0. Last but not least, the worst case: *δ* = 0 and *δ*_*S*_ > 0, meaning that *F*_*S*_ females re-mate but not *F*_*W*_ females, then

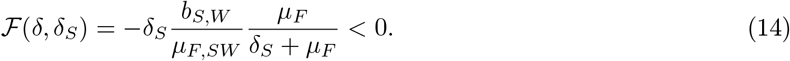

In that case, re-mating *F*_*S*_ females only, will considerably and negatively impact SIT. We will illustrate the different cases in section 3.1, page 14.

In the next proposition, where 0 *≥ ε ≥ ε*_max_, we show existence of a critical threshold, 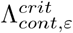, for the release rate, Λ, above which no positive steady state can exists.

#### Proposition 1

*Assume ℛ* > 1 *then the following results hold true for system* (2):

- *Assume ε* > *ε*_max_, *then there always exists one positive steady state* **E**^*^ >> 0, *whatever the size of the sterile male releases*.
- *Assume ε* = *ε*_max_, *then, setting* 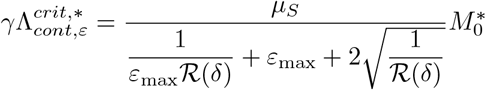, *we deduce that*
  – *If*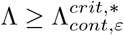, *there is no positive steady state*.
  – *If*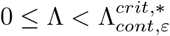, *there is one positive steady state* 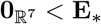
- *Assume* 0 *≤ ε* < *ε*_max_, *then, there exists* 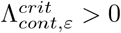 *such that*
  – *If* 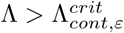, *there is no positive steady state*.
  – *If* 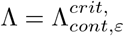, *there is one positive steady state*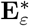.
  – *If* 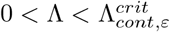, *then there are two positive steady states* **E**_*ε,−*_ *and* **E**_*ε*,+_, *such that*

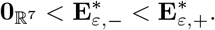

**Proof**. See Appendix C, page 33.

This proposition, combined with Lemma 2, is fundamental to show that pest control, and eventually elimination, is possible as long as we release a sufficiently la rge am ount of sterile ma les, i. e.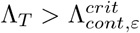.

However, lemma 2 ensures elimination only if the wild population is sufficiently small. For practical applications, and, in particular, to ensure that elimination is still possible when the wild population is large, we need to extend the result of Lemma 2 and show that 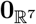 is GAS. This is done using the auxiliary system (2)

#### Theorem 2

*Assume ε* < *ε*_max_ *and* 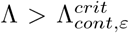, *then* 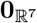 *is GAS for the monotone auxiliary system* (3), *and thus GAS for system* (2).

**Proof**. See appendix D, page 36.

#### Remark 4

*According to Theorem 2, provided that ε is sufficiently small, it is possible to reach (asymptotically) elimination when* Λ *is large enough*.

We can deduce from Proposition 1 and Theorem 2, that the residual fertility, *ε*, is an essential parameter to take into account when designing SIT programs. As *ε* increases, the SIT becomes less effective and, eventually, fails to reduce the wild population.

The precision and homogeneity of the sterilization step is very important. Although the irradiation process is very well mastered, the level of residual fertility will depend on the homogeneity of the dose delivered to the fly pupae, which may be affected by container volume, irradiation equipment and source.

The classical recommendation is to have the lowest residual fertility possible. Here we provide, for the first time, insights on the impact of RF on the release success.

Technical and production improvements in medfly SIT have been obtained using genetic sex strains [52] in which males also have a naturally reduced fertility level (49.29%).The residual fertility of the VIENNA-8 GSS strain was reported to 0.84% ± 0.08% (0.6% ± 0.13%) at 100*Gy* (120*Gy*) [52][Table 3] on average. However, a higher residual fertility was observed for a similar strain (VIENNA-8 GSS strain with a Valencian background) irradiated at the same dose but with an electron beam accelerator [5], i.e.1.13% ± 0.55% for 100Gy.

Another important aspect is the re-mating of females: having good knowledge of this phenomenon can significantly change the constraint on the residual fertility, *ε*. We will illustrate the impact of re-mating in the numerical section.

However, it is important to have in mind that targeted crops may also affect the effectiveness of a SIT program. Indeed, Ceratitis capitata’s basic reproduction numbers depends on the main fruit hosts and, thus, can take a large range of values. For instance, according to [45], estimates of ℛ_0_ on deciduous fruits, like nectarine (or plum), yield _0_ *≈* 227.28± 89.1 (ℛ_0_ *≈* 276.59 ± 54.15), at 25^*o*^C, whereas ℛ_0_ *≈* 27.05 on citrus, at 24^*o*^C [53]. Clearly the constraint on *ε* will depend on the targeted host crop: for instance, a residual fertility of 2% might be acceptable for citrus but not necessarily acceptable for nectarine or plum: compare 1/27.05 *≈* 0.037 and 1/227.05*≈* 0.0044. Section 5, where we apply our results to a real crop, we will consider medfly data related to peach, obtained in Tunisia [44], as Tunisia and Corsica show similarities from the agricultural context point of view.

#### Remark 5

*Obtaining theoretically the stability properties of the positive equilibria*, 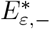, *and* 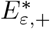, *when they exist, is not easy considering the complexity of the system. The numerical simulations indicate that* 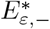 *is unstable while* 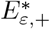 *is asymptotically stable, and that the equilibria are the only invariant set of the system on* 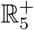. *However, we provide in appendix C, conditions to verify, at least numerically, if they are unstable or not*.

### 2.3 Periodic impulsive SIT releases

We assume now that sterile males are released periodically, every *τ* days. Assuming each release as an instantaneous event, this situation can be modeled using a semi-discrete approach, like in [51, 16, 21]. Thus, model (1) becomes

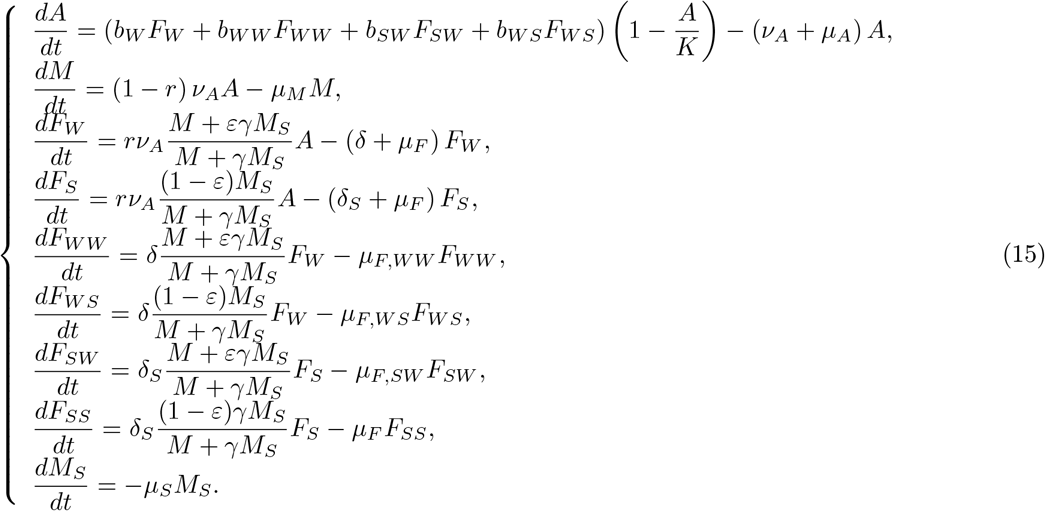

with

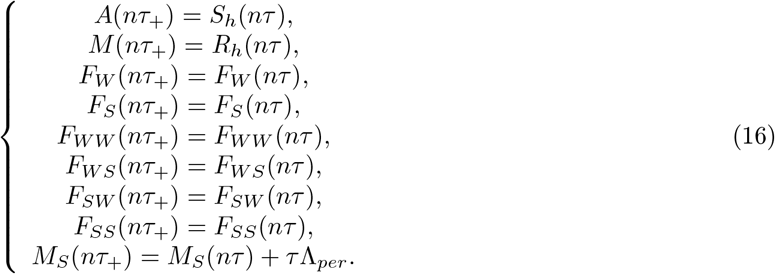

It is straightforward to show that, as *t →* +*∞, M*_*S*_ converge toward the periodic solution

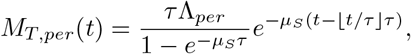

such that system (15)-(16) can be reduced to

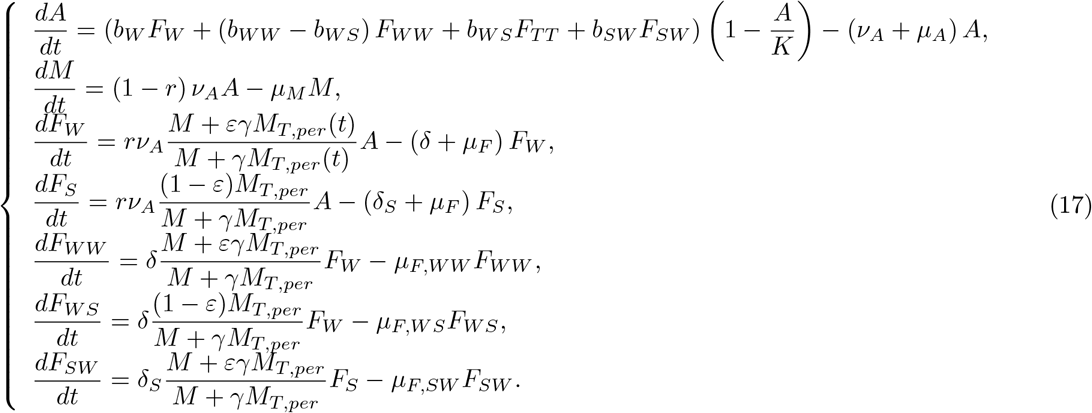

Setting

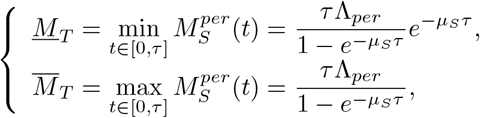

it is obvious to deduce that, for *t* sufficiently large, system (17) is lower and upper bounded by the following two monotone systems

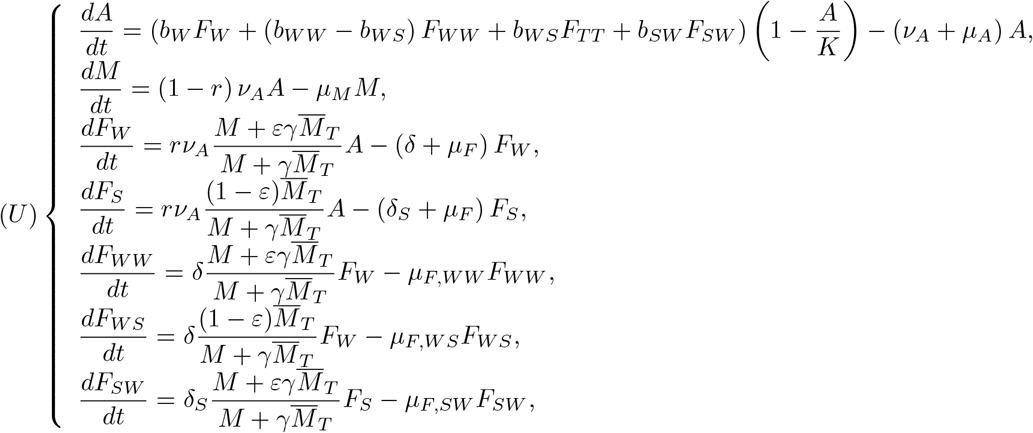

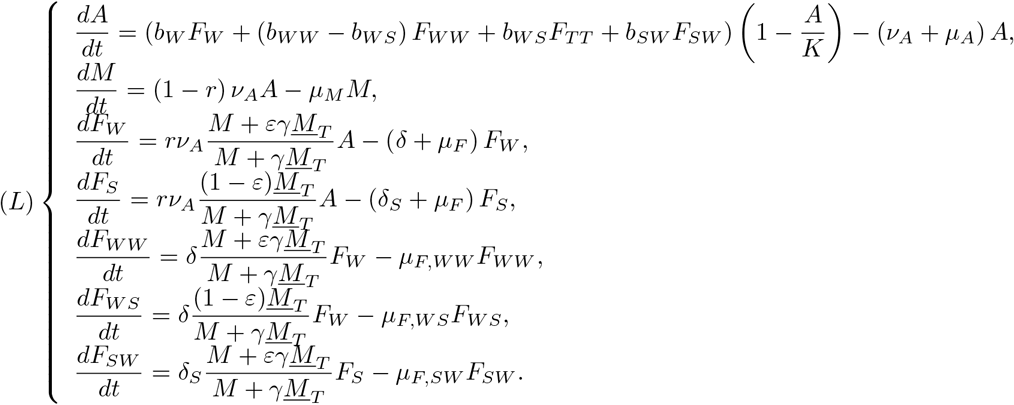

Then, applying Proposition 1 to system (L) or system (U), we obtain

#### Proposition 2

*Assume ℛ* > 1 *and ε* < *ε*_max_.

- *When* 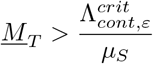, *the trivial equilibrium* 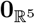 *is Globally Asymptotically Stable for system (U)*.
- *When* 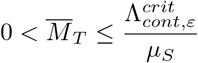, *system (L) admits one or two positive equilibria Ē*_1,5*D*_ *≤ Ē*_2,5*D*_. *In addition, if the initial data of (L) is greater than or equal to E*_1,5*D*_, *then the corresponding solution is also greater than or equal to E*_1,5*D*_. *Similarly, since* 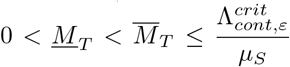, *then system (U) admits one or two positive equilibria* 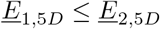. *Finally, the set* 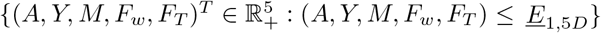 *belongs to the basin of attraction of* 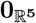 *for system (U), hence by comparison, it belongs to the basin of attraction of* 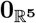 *for system* (17).

From the previous remarks, we can deduce

#### Proposition 3

*Assume ℛ* > 1, *ε* < *ε*_max_, *and*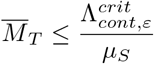, *the set*

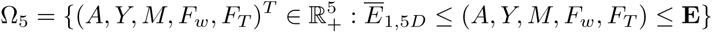

*is positively invariant by system* (17), *where* **E**, *the initial wild equilibrium, is defined in Theorem 1*.

Finally, using the previous result and Brouwer fixed point theorem, with comparison arguments, it is possible to show

#### Theorem 3

*Assume ℛ*> 1, *ε* < *ε, and*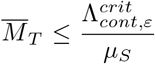. *Then, for each initial condition in* Ω_5_, *system* (17) *has at least one positive τ -periodic solution* **E**_*per*_ *such that* **E**_*per*_ ∈ Ω_5_.

The previous results are obtained by comparison arguments using systems (L) and (U). While they ensure the existence of periodic critical threshold, 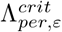, it is not possible to find an analytical formula, like for the continuous case. However, it is possible to estimate 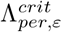 numerically, by solving system (17), page 11, using an iterative approach.

## 3 Results

We now apply the previous theoretical result to medfly, *Ceratis capitata*, parameters. We summarize in table 2, page 13, the values estimated for some biological parameters obtained from the literature for medfly grown on peach (when available) and on sterile males from the V8 GSS strain (Vienna-8 Genetic Sexing Strain).

**Table 2:**
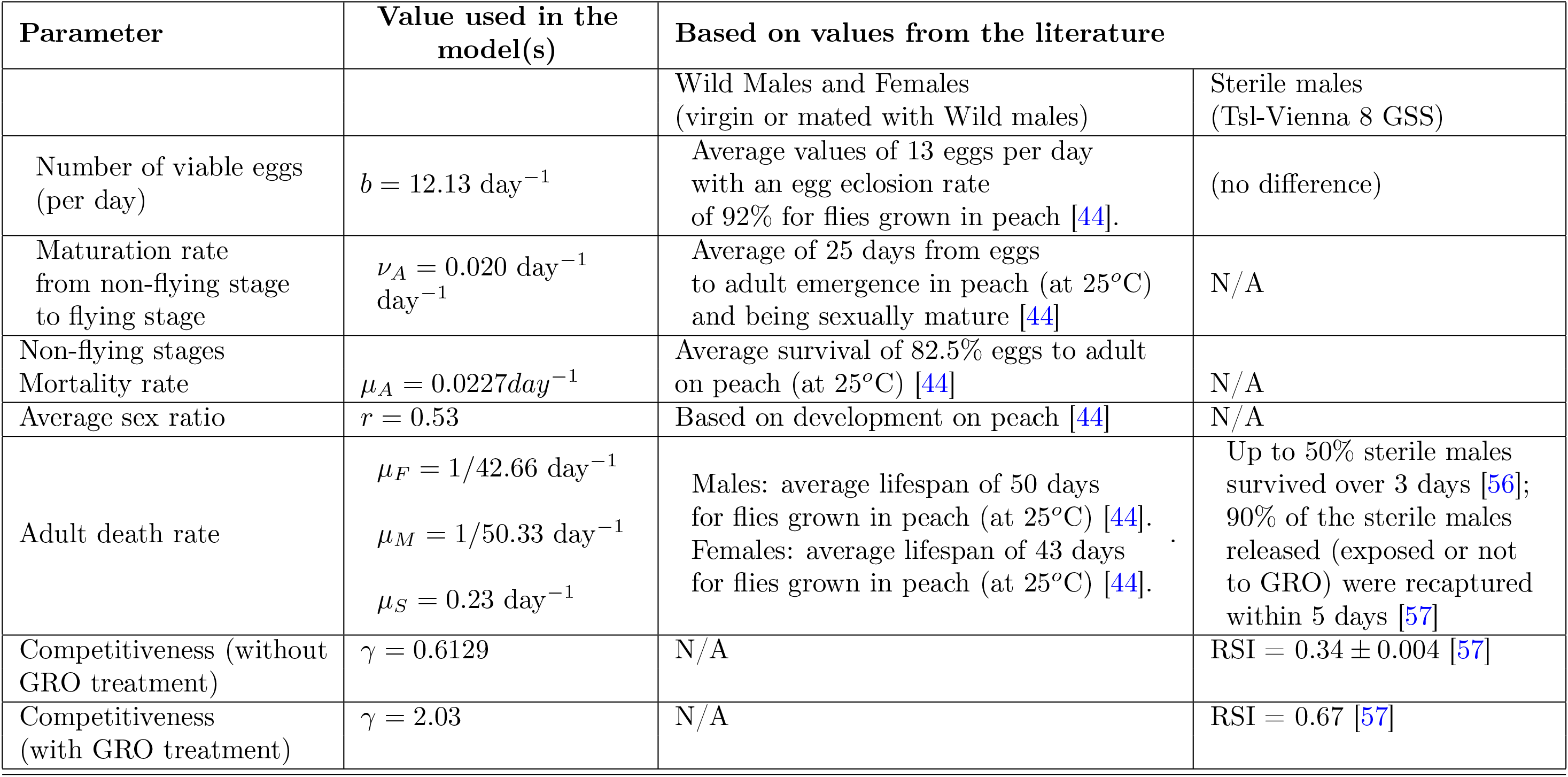
*C. capitata* entomological parameter values used in this model (literature selected for demographic parameters were studies using host fruits rather than artificial diet, and field studies when available). The parameters values for *δ* and *δ*_*S*_ are given below in Table 3

| Parameter | Value used in the model(s) | Based on values from the literature |  |
| --- | --- | --- | --- |
|  |  | Wild Males and Females (virgin or mated with Wild males) | Sterile males (Tsl-Vienna 8 GSS) |
| Number of viable eggs (per day) | $b = 12.13 \text{ day}^{-1}$ | Average values of 13 eggs per day with an egg eclosion rate of 92% for flies grown in peach [44]. | (no difference) |
| Maturation rate from non-flying stage to flying stage | $\nu_A = 0.020 \text{ day}^{-1}$ | Average of 25 days from eggs to adult emergence in peach (at 25°C) and being sexually mature [44] | N/A |
| Non-flying stages Mortality rate | $\mu_A = 0.0227 \text{ day}^{-1}$ | Average survival of 82.5% eggs to adult on peach (at 25°C) [44] | N/A |
| Average sex ratio | $r = 0.53$ | Based on development on peach [44] | N/A |
| Adult death rate | $\mu_F = 1/42.66 \text{ day}^{-1}$<br>$\mu_M = 1/50.33 \text{ day}^{-1}$<br>$\mu_S = 0.23 \text{ day}^{-1}$ | Males: average lifespan of 50 days for flies grown in peach (at 25°C) [44].<br>Females: average lifespan of 43 days for flies grown in peach (at 25°C) [44]. | Up to 50% sterile males survived over 3 days [56]; 90% of the sterile males released (exposed or not to GRO) were recaptured within 5 days [57] |
| Competitiveness (without GRO treatment) | $\gamma = 0.6129$ | N/A | RSI = $0.34 \pm 0.004$ [57] |
| Competitiveness (with GRO treatment) | $\gamma = 2.03$ | N/A | RSI = 0.67 [57] |

All computations have been done using codes developed with the software R-studio [54] and R [55]. In particular, estimates of the critical periodic ratio,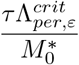, have been obtained by solving system (15), page 10, using a nonstandard finite difference scheme (see [18] and references therein), in order to derive a fast algorithm to obtain each figure in a reasonable amount of time, on a laptop.

In the forthcoming simulations, we derive estimates of the critical thresholds, 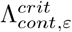 and 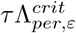 over the wild males at equilibrium, 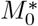, because this is how releases densities are defined in SIT programmes.

### 3.1 Parameterization and simulations

The difficulty of using modeling to understand biological phenomena lies in the estimate of the values of the biological parameters. Most behavioral or biological studies are made in laboratory settings and under controlled conditions; they do not necessarily reflect behavior in the field. On the other hand, reports from field studies may cover only a small portion of the diversity of behaviors or physiology of wild populations. Most of the parameters values here were estimated directly or indirectly from the literature, selecting studies that brought the closest estimates to what may occur in the field rather than laboratory studies, when possible.

Since we have only part of the whole parameters, we will “adapt” the parameters values from [44], on peach, and also use results from [37, 36, 40] for some parameters related to re-mating.

We estimated *ν*_*A*_ and *μ*_*A*_ from the data provided on peach development [44] as follows. Since 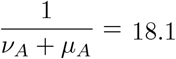 [44][Table 2], and, the proportion of hatched eggs that become adults is 66.09% [44][Table 3], this means that 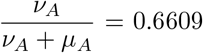, that is 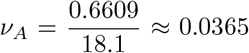 and *μ*_*A*_ = 0.0187. Similarly, we deduce that *b*_*W*_ = 13.19 × 0.92 = 12.135 [44][Tables 1 and 3].

Little literature exists on development parameters from fruit host rather than artificial diet. Big variations can occur according to the host, citrus or clementine having lower developmental rate and longer duration but a higher adult lifespan, as compared to peach hosts [44].

The model from Plant & Cunningham [56] predicted that 50% of the released sterile males were dead after 3 days; assuming that the population size is given by 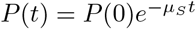 we derive that 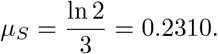.

The estimation of the sterile male competitiveness index is based on the Relative Sterility Index (RSI), through cage experiments. Without GRO treatment, the mean RSI is low, around 0.38, but still greater than the threshold 0.2, leading to a competitiveness parameter *γ* = 0.61 [48]. When sterile males are treated with an optimal dose of ginger root oil (GRO) [48], then the mean RSI becomes 0.67 which implies a competitive parameter *γ* = 2.03. Note that sterile and wild males are equally competitive when RSI= 0.5, for a 1 : 1 : 1 density. The parameters values are summarized in Table 2, page 13.

Some field studies reported percentages of double mating in wild females: up to 50% [28]; < 28% [29]; 4 to 28% [58]. Laboratory tests reported by Abraham et al [25] indicated 20% re-mating for females mated with 100Gy-irradiated males. Recently, Pogue et al (2022)have shown that female mating propensity reduces over time [59]: with an average of 75% females *C*.*capitata* re-mating a second time but only less than 25% performing successive re-mating. The litterature on medfly remating varies with studies and the global picture is not yet complete, as reviewed by Pérez-Staple & Abraham [60]. Here, we’ve chosen to estimate the parameters *δ* and *δ*_*S*_ based on the values reported from laboratory experiments by [30, Table 1]. From competing scenarios with wild and sterile males, we could estimate that the re-mating rate was equal to the proportion of re-mating, (40% when females are mated first with a wild male), divided by the mean refractory period (RFP), (i.e. 2.5 days), such that we get 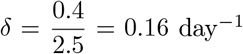 for females exposed to sterile males not treated with GRO. Using the same reasoning we also derive *δ*_*S*_ with and without GRO-treatment: The lifespan of sterile males was reported not to be impacted by the GRO treatment (see [57]).

Re-mating has an important impact on females fitness: see [38, 40, 36]. In [40], single-mated females live on average 27 days, while multiple-mated females live on average 34 days, such that we deduce 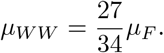.

Similarly, the average number of offspring (not the amount of (hatched) eggs deposit every day) is, on average, 87 for single-mated females, and 142 for double-mated female over their lifetime. Thus daily, single-mated females produces, on average, 3.22 offspring per day, and multiple-mated females produces 4, 1765 offspring per day. Thus, we deduce that the production of (hatched) eggs for a double-mated female is, on average,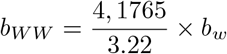. This last result, *b*_*W W*_ > *b*_*W*_, is confirmed in [38].

For the females, *F*_*W W*_, *F*_*W S*_ and *F*_*SW*_, there is no fecundity and lifespan data available related to remating with the V8-strain. Only few contrasting (partial) data are available for *C. capitata* in [36] and [37], from which we can extrapolate some numerical values of *b*_*W S*_, *b*_*SW*_, but not for the death-rates, *μ*_*W S*_, and *μ*_*SW*_. In [36], the authors considered two genotypes: they showed that copulation order with different genotypes (irradiated or not) may influence the fitness of the double-mated females. We will consider [37], even if some data are missing. Using Table 2 [37], we derive 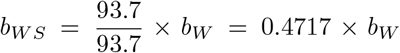 and 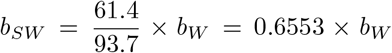. These are of course rough estimate. It would be preferable to have experiments estimating simultaneously all parameters *b*_*_ and *μ*_*_. Until then, we will assume that *μ*_*W S*_ = *μ*_*SW*_ = *μ*_*F*_.

#### Remark 6

*It is interesting to notice that data to estimate b*_*W S*_ *and b*_*SW*_ *also exist for B. dorsalis in [61], showing the first male sperm precedence. Indeed, if the total amount of laid eggs is (statistically) similar for F*_*W S*_ *and F*_*SW*_, *the proportion of eggs hatched is not: it is* 71.7% *for F*_*W S*_ *and* 54.9% *for F*_*SW*_. *It is important to notice that this result shows that the first sperm seems to have the precedence to the second sperms since b*_*W S*_ > *b*_*SW*_, *at least for B. dorsalis. There is no information about the death-rates, μ*_*W S*_ *and μ*_*SW*_.

From [62], for *C. capitata*, there is a tendency of second sperm precedence, at least for the first ovipositions, and then it decreases in favor of the first sperm. However, it would be more than welcome to conduct experiments, like [61], to clearly estimate the total amount of hatching eggs laid by *F*_*W S*_, *F*_*SW*_ and, also *F*_*W W*_, and also their mean lifespan, to estimate *b*_*_ and *μ*_*_.

Using Table 3, we derive Table 4, page 15, where the values for the basic offspring number are computed according to the two different sub-cases: with re-mating, ℛ (*δ*), without re-mating, ℛ (0), with and without GRO-treatment. As shown, the impact of re-mating is important on the basic offspring number: the smaller the refractory period the larger the basic offspring number. Clearly, with or without GRO-treatment, there is a need to study the mating and re-mating behaviour of females that mated either with wild males or sterile males.

**Table 3:** Re-mating rates with and without GRO treatment [30].

| | $\delta$ | $\delta_S$ |
| --- | --- | --- |
| SM | $0.4/2.5 = 0.16$ | $0.64/2.02 = 0.3161$ |
| SM-GRO | $0.2/2 = 0.1$ | $0.34/1.44 = 0.2361$ |

**Table 4:**
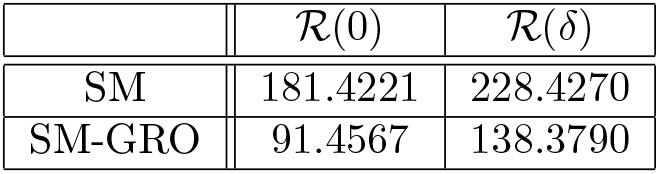
Basic Offspring numbers with and without GRO treatment according to Tables 2 and 3.

The tolerable value for residual fertility is very low according to the chosen parameters values. This would give reason to SIT implementation programs that choose a fully sterilizing dose, such as [25].

Solving equation (22), page 34, we first derive estimates of 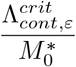 in order to highlight the issues of residual fertility and re-mating in SIT control treatment. In the forthcoming simulations we consider four cases for the re-mating rates:

a. Assume that *δ* = 0 and *δ*_*S*_ > 0. This means that female that mated with a sterile male will always re-mate. This, would be the worst case, but it is interesting to see how this impacts the SIT treatment.
b. No re-mating, i.e. *δ* = *δ*_*S*_ = 0: females mate only once whatever males are wild or sterile.
c. With re-mating, such that *δ* = *δ*_*S*_ > 0. This is what we called “equal remating”, i.e. female will re-mate at the same rate, whatever if they mated with a wild or a sterile males.
d. With re-mating, such that 0 < *δ* < *δ*_*S*_. This is supposed to be the “standard” case for *C. capitata*: after mating with a sterile male, a female will re-mate faster than a female that mated with a wild male.

We also consider different numerical values for *b*_*W S*_ and *b*_*SW*_ for continuous and periodic releases to illustrate the importance of the double-mated females parameters. In all simulations, SIT-treatment start at time *t* = 100 days.

Case 1. Assume *b*_*W,S*_ = *b*_*S,W*_ = 0.5 × *b*_*W*_. From Fig. 2, page 16, GRO-treatment improves drastically the ratio between the critical release rate and the amount of wild males at equilibrium,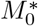: see Fig. 2, page 16. The gain in release rate is nearly 10 with the GRO treatment compared to without the GRO treatment. The larger the residual fertility, the larger the amount of sterile males to release. However, it is interesting to notice that this increase is low for *ε* < 0.25%. Of course, as expected, the worst case is the case when matings with sterile males induce a re-mating, *δ*_*S*_ > 0, while there is no re-mating when matings occur with wild male, *δ* = 0 (the blue curve). From Table 5, we see that GRO-treatment does not change much the upper bound for *ε*_*max*_.

**Table 5:** Numerical estimates for *ε*_max_ with and without GRO treatment - Case 1: *b*_*W,S*_ = *b*_*S,W*_ = 0.5 × *b*_*W*_.

| | $\delta_S > \delta > 0$ | $\delta_S = \delta > 0$ | $\delta_S > 0, \delta = 0$ | $\delta_S = \delta = 0$ |
| --- | --- | --- | --- | --- |
| $\varepsilon_{\max}$ | 0.00534 | 0.00549 | 0.00376 | 0.00551 |
| $\varepsilon_{\max, GRO}$ | 0.00524 | 0.00549 | 0.00379 | 0.00551 |

**Figure 2:**
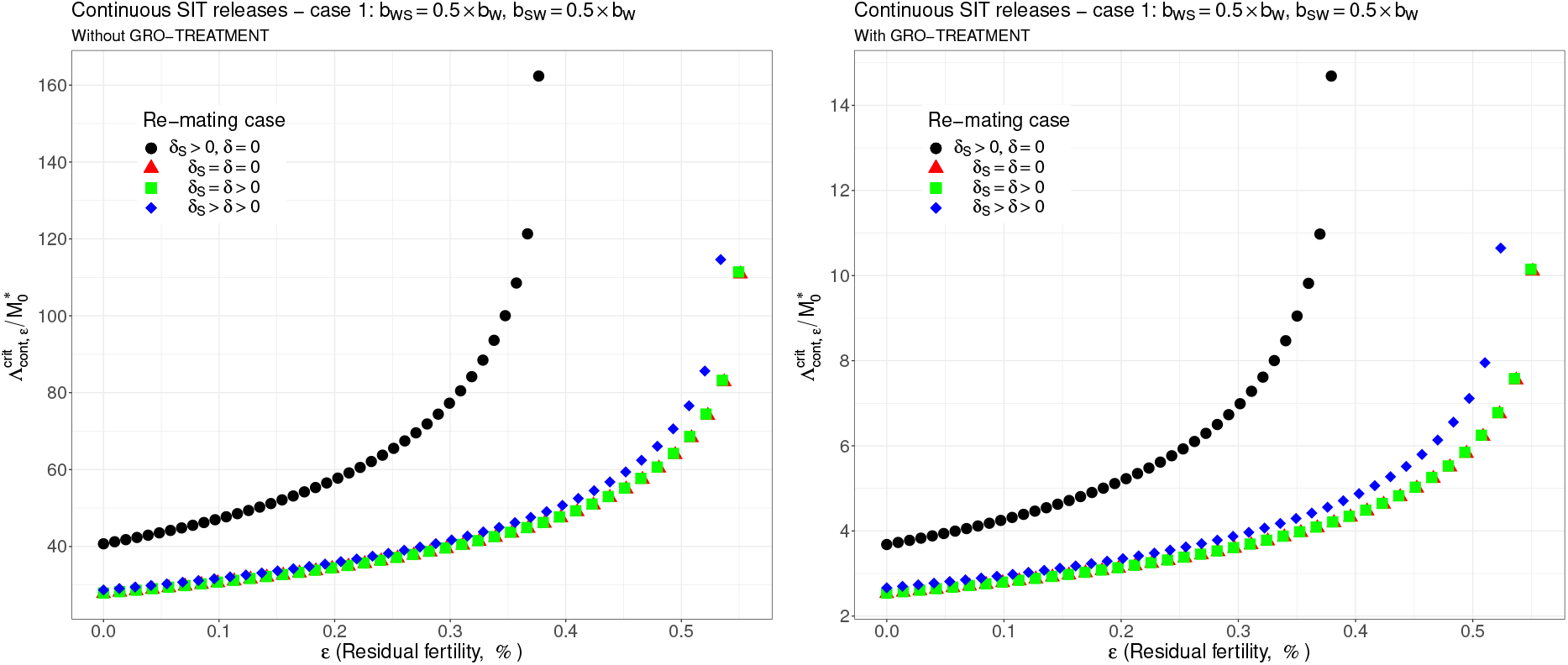
Critical ratio for continuous releases as a function of residual fertility - re-mating case 1 with *b*_*W,S*_ = 0.5 × *b*_*W*_, *b*_*S,W*_ = 0.5 × *b*_*W*_ : (a) without GRO-treatment (b) With GRO-treatment. Simulations with different re-mating configurations: the black bullets, with re-mating rates *δ*_*S*_ > 0 and *δ* = 0; the red triangles, the NO re-mating case, *δ*_*S*_ = *δ* = 0; the green squares, with positive and equal re-mating rates, *δ*_*S*_ = *δ* > 0; the blue diamonds, with positive re-mating rates, *δ*_*S*_ > *δ* > 0.

Case 2. As explained above, and following [37], we assume now that *b*_*W S*_ = 0.4717 × *b*_*W*_ and *b*_*SW*_ = 0.6553× *b*_*W*_ : see Fig. 3, page 17). Contrary to case 1, re-mating mainly impacts *ε*_max_: see Table 6, page 16. Indeed,

**Table 6:** Numerical estimates for *ε*_max_ with and without GRO treatment - Case 2: *b*_*W S*_ = 0.4717 × *b*_*W*_ and *b*_*SW*_ = 0.6553 × *b*_*W*_.

| | $\delta_S > \delta > 0$ | $\delta_S = \delta > 0$ | $\delta_S > 0, \delta = 0$ | $\delta_S = \delta = 0$ |
| --- | --- | --- | --- | --- |
| $\varepsilon_{\max}$ | 0.00479 | 0.00495 | 0.00343 | 0.00551 |
| $\varepsilon_{\max, GRO}$ | 0.00471 | 0.00499 | 0.00345 | 0.00551 |

**Figure 3:**
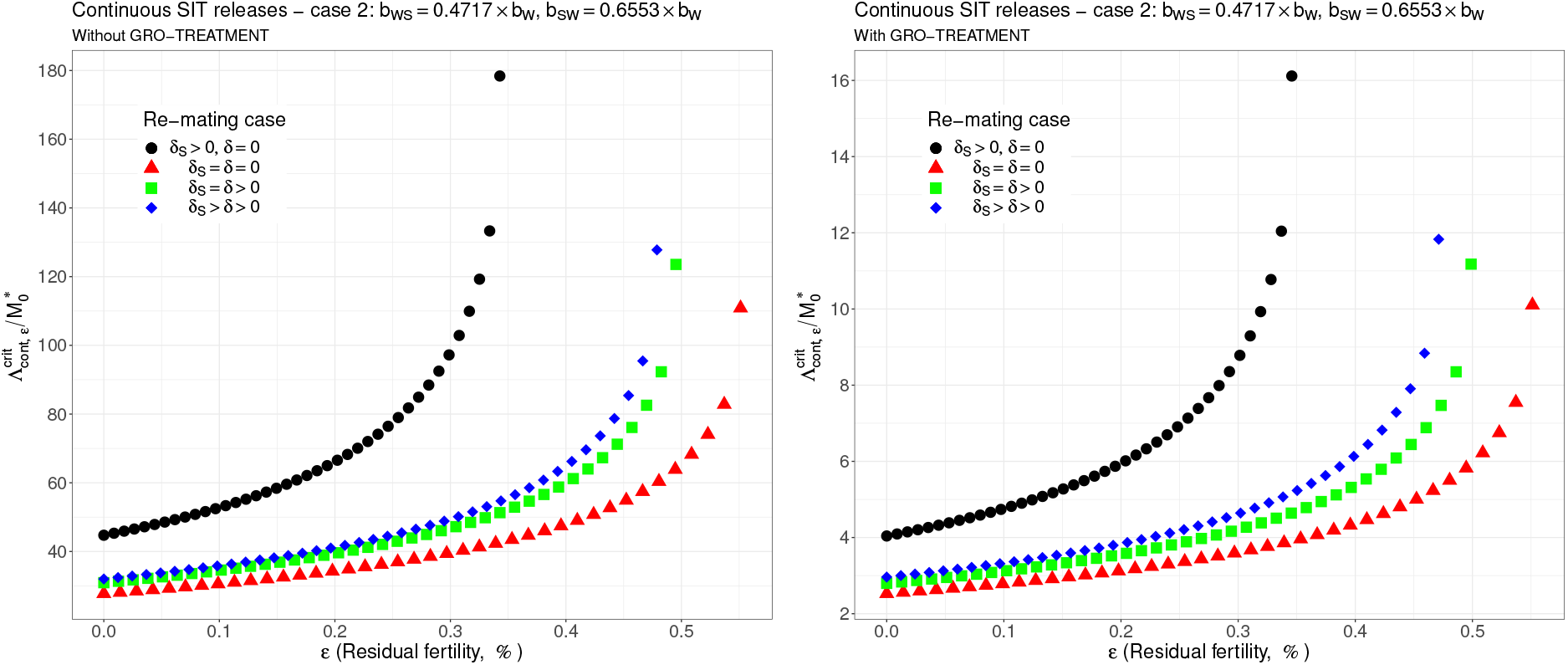
Critical ratio for continuous releases as a function of residual fertility - re-mating case 2 with *b*_*W,S*_ = 0.4717 × *b*_*W*_ and *b*_*SW*_ = 0.6553 × *b*_*W*_ : (a) without GRO-treatment (b) With GRO-treatment. Simulations with different re-mating configurations: the black bullets, with re-mating rates *δ*_*S*_ > 0 and *δ* = 0; the red triangles, the NO re-mating case, *δ*_*S*_ = *δ* = 0; the green squares, with positive and equal re-mating rates, *δ*_*S*_ = *δ* > 0; the blue diamonds, with positive re-mating rates, *δ*_*S*_ > *δ* > 0.

when re-mating occurs, with *δ*_*S*_ > *δ* > 0, the maximal residual fertility decreases, *ε*_max_ = 0.0047, compared to case 1, where *ε*_max_ = 0.0052. However, the values for the critical release rate are similar between case 1 and case 2, at least when *ε* < 0.25%. The GRO-treatment does not change much the upper bound for *ε*_*max*_.

Case 3. This case is more or less related to the results obtained in [36]: see cases F and G in Table 2, where the sterile male is not of the same genotype than the wild male. Such a case could eventually occur. Thanks to Tables 1 and 2 in [36], we assume that *b*_*W S*_ = 0.1532 × *b*_*W*_ and *b*_*SW*_ = 0.65 × *b*_*W*_ : see Fig. 4, page 18. As expected, this is a case where re-mating is favorable for SIT: *ε*_max_ takes larger values when *δ* > 0: see Table 7, page 17. Note also, that in [36][Table 3], the cases E and H (the reverse cases to cases F and G) may be even more favorable for re-mating. However, there is no change in the critical release rates compared to cases 1 and 2, except for the re-mating cases where *δ*_*S*_ *≥ δ* > 0: see Fig. 4, page 18.

**Table 7:** Numerical estimates for *ε*_max_ with and without GRO treatment - Case 3: *b*_*W S*_ = 0.1532 × *b*_*W*_ and *b*_*SW*_ = 0.65 × *b*_*W*_.

| | $\delta_S > \delta > 0$ | $\delta_S = \delta > 0$ | $\delta_S > 0, \delta = 0$ | $\delta_S = \delta = 0$ |
| --- | --- | --- | --- | --- |
| $\varepsilon_{\max}$ | 0.00629 | 0.00661 | 0.00344 | 0.00551 |
| $\varepsilon_{\max, GRO}$ | 0.00607 | 0.00652 | 0.00347 | 0.00551 |

**Figure 4:**
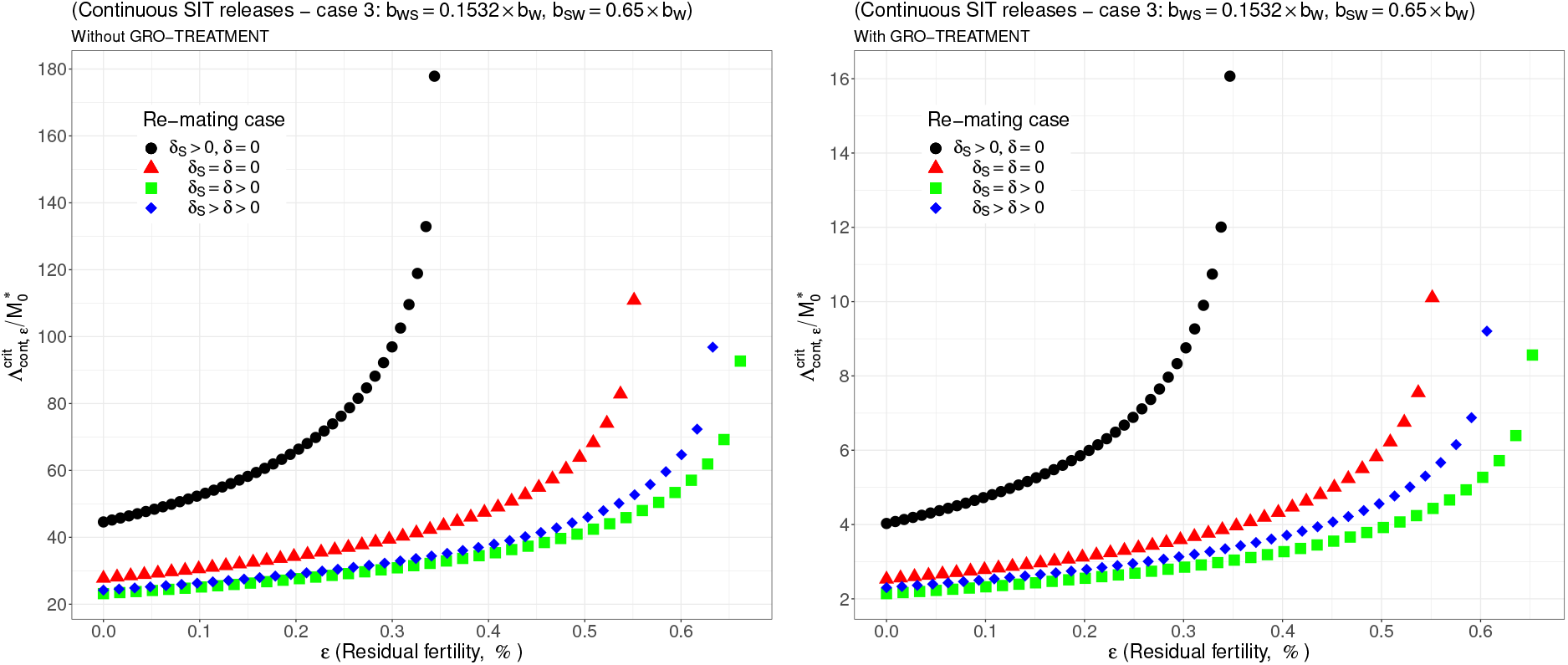
Critical ratio for continuous releases as a function of residual fertility - re-mating case 3 with *b*_*W S*_ = 0.1532 × *b*_*W*_ and *b*_*SW*_ = 0.65 × *b*_*W*_ : (a) without GRO-treatment (b) With GRO-treatment. Simulations with different re-mating configurations: the black bullets, with re-mating rates *δ*_*S*_ > 0 and *δ* = 0; the red triangles, the NO re-mating case, *δ*_*S*_ = *δ* = 0; the green squares, with positive and equal re-mating rates, *δ*_*S*_ = *δ* > 0; the blue diamonds, with positive re-mating rates, *δ*_*S*_ > *δ* > 0.

Case 4. The worst case: re-mated Females *F*_*W,S*_ and *F*_*S,W*_ are always using the fertile sperms (sterile sperm is not competitive or females are able to distinguish between sterile and fertile sperms): *b*_*W,S*_ = *b*_*S,W*_ = *b*_*W*_. The results are almost similar for all re-mating cases: see Fig. 5, page 18.

**Figure 5:**
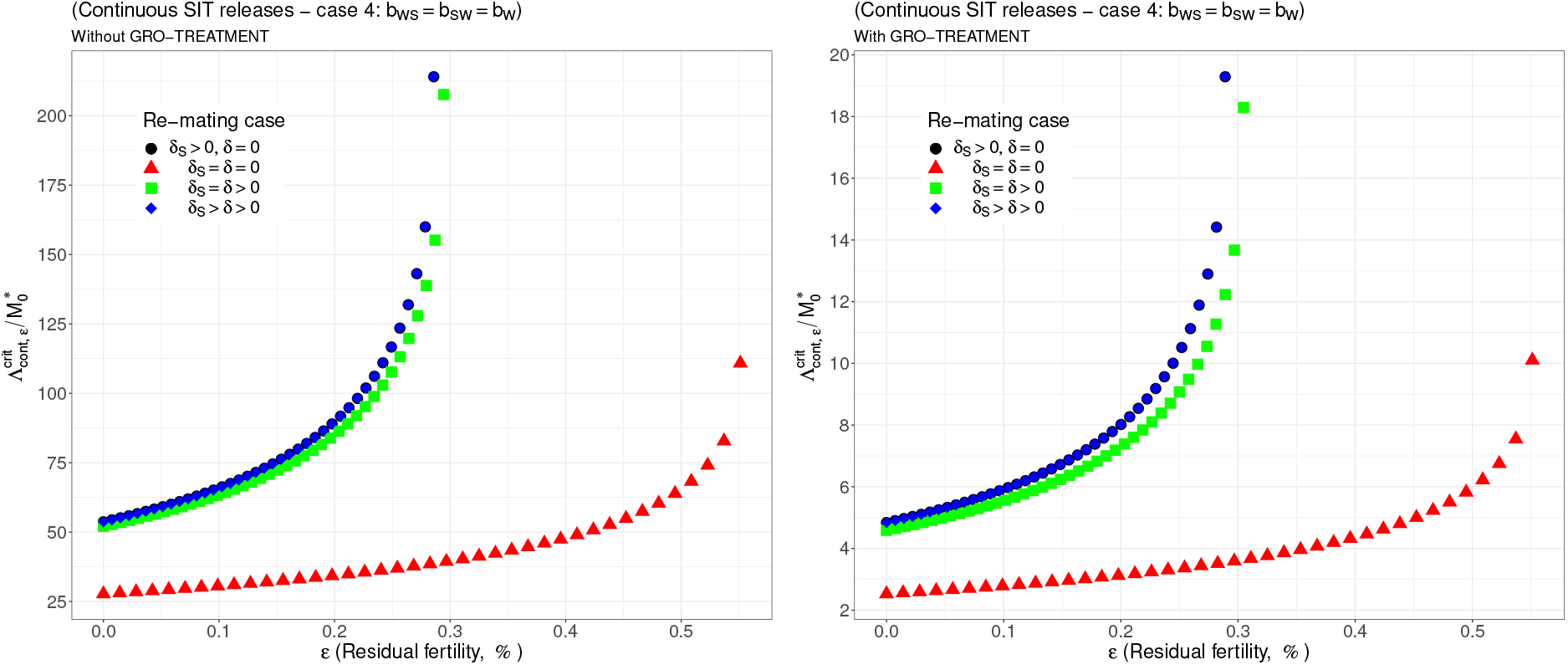
Critical ratio for continuous releases as a function of residual fertility - re-mating case 4 with *b*_*W,S*_ = *b*_*W*_, *b*_*S,W*_ = *b*_*W*_ : (a) without GRO-treatment (b) With GRO-treatment. Simulations with different re-mating configurations: the black bullets, with re-mating rates *δ*_*S*_ > 0 and *δ* = 0; the red triangles, the NO re-mating case, *δ*_*S*_ = *δ* = 0; the green squares, with positive and equal re-mating rates, *δ*_*S*_ = *δ* > 0; the blue diamonds, with positive re-mating rates, *δ*_*S*_ > *δ* > 0.

We can notice that in all simulations, the gain, in the critical release ratio, is almost of factor 10 with the GRO-treatment i. On the other hand there is almost no gain on *ε*_max_.

It is interesting to compare the upper bounds, *ε*_max_, given in Tables 5-8, with real values. For instance, in [52][Table 3], the sterility induced by 100GY (120GY) sterilized males was 99.42% ± 0.1 (99.72% ± 0.15), that is an average residual fertility of 0.58% (0.28%), when considering the percentage of pupae surviving from progeny of females mated by sterilized males. This is close to the upper bounds given in Tables 5 and 7, whatever re-mating occurs, with *δ* > 0, or not. On the other hand, in [5], the sterility induced by 100GY sterilized males was 98.87%± 0.55 providing an average residual fertility of 1.13%, that is *ε* = 0.0113. This value is much larger than the values given in Tables 5, 6 and 8, but lower than the values given in Table 7

**Table 8:** Numerical estimates for *ε*_max_ with and without GRO treatment - Case 4: *b*_*W,S*_ = *b*_*S,W*_ = *b*_*W*_.

| | $\delta_S > \delta > 0$ | $\delta_S = \delta > 0$ | $\delta_S > 0, \delta = 0$ | $\delta_S = \delta = 0$ |
| --- | --- | --- | --- | --- |
| $\varepsilon_{\max}$ | 0.00285 | 0.00294 | 0.00286 | 0.00551 |
| $\varepsilon_{\max, GRO}$ | 0.00289 | 0.00304 | 0.00290 | 0.00551 |

when re-mating occurs. Following our theoretical results, this means that the sterile males released under the conditions of [5] could only induce a reduction of the wild population, but not an elimination, at least in cases 1, 2 and 4. In fact, it seems clear that estimating the residual fertility alone is not sufficient: the impact of RF might depend on the values taken by *b*_*_, *μ*_*_, *δ, δ*_*S*_, and also *γ*.

For periodic releases, we fully rely on numerical simulations of system (17), page 11, to estimate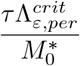, for a given period of releases, *τ*. We consider the four cases studied in the continuous release part. As expected, whatever the case, the weekly releases provide bad results, i.e. very massive releases are requested: see from Fig. 6, page 19, to Fig. 9, page 20. This is not surprising as we considered a short lifespan for sterile males. The best release period is 3 days as showed from Fig. 10, page 21, to Fig. 13, page 22. Again, the GRO-treatment is very helpful, and if residual fertility is low, say less than 0.3%, then the release ratios are reasonable in terms of production: see all left figures from Fig. 6, page 19, to Fig. 13, page 22. Note also the critical release ratio for periodic releases cannot directly be deduced from the continuous critical release ratio, Λ^*crit*^, by multiplying it by *τ*, the release period, because, in general,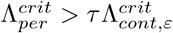.

**Figure 6:**
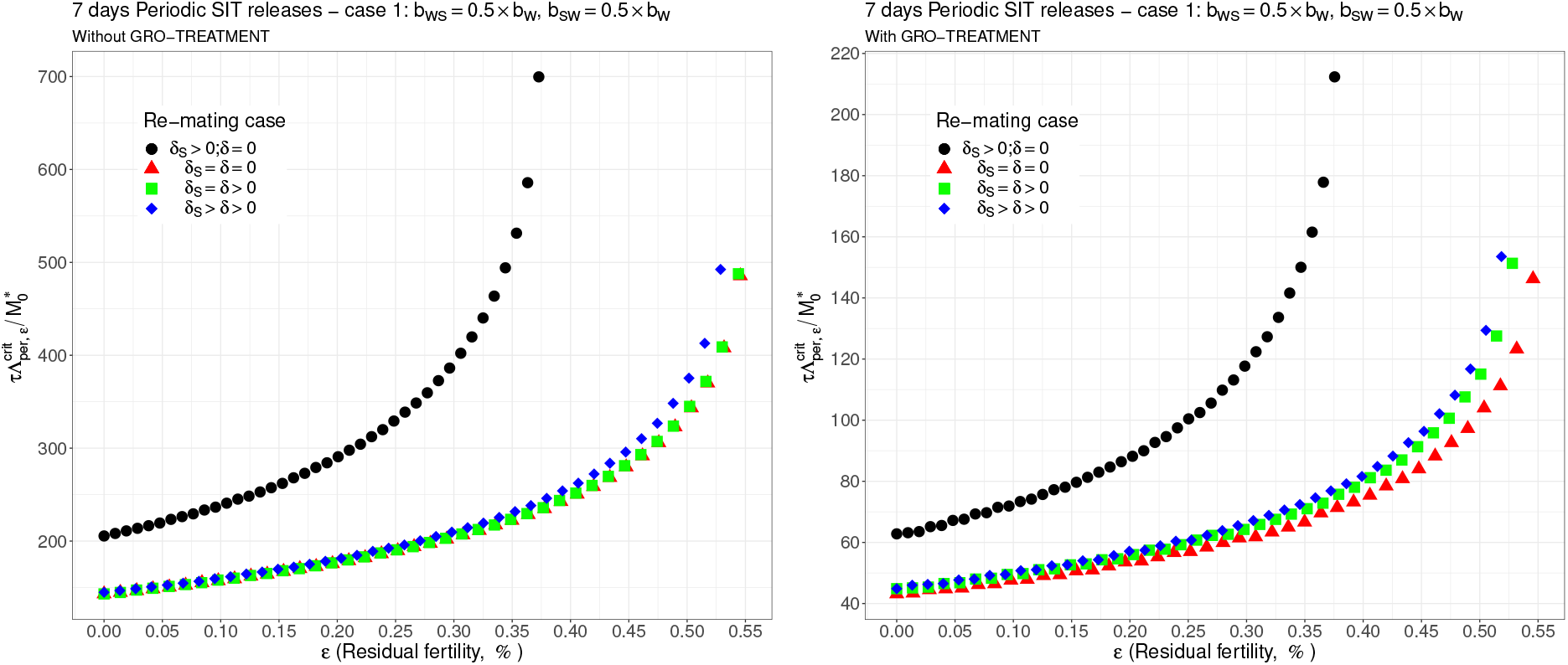
Critical ratio for weekly periodic releases as a function of residual fertility - re-mating 1 with *b*_*W,S*_ = 0.5 ×*b*_*W*_, *b*_*S,W*_ = 0.5 ×*b*_*W*_. Simulations with different re-mating configurations: the black bullets, with re-mating rates *δ*_*S*_ > 0 and *δ* = 0; the red triangles, the NO re-mating case, *δ*_*S*_ = *δ* = 0; the green squares, with positive and equal re-mating rates, *δ*_*S*_ = *δ* > 0; the blue diamonds, with positive re-mating rates, *δ*_*S*_ > *δ* > 0.

**Figure 7:**
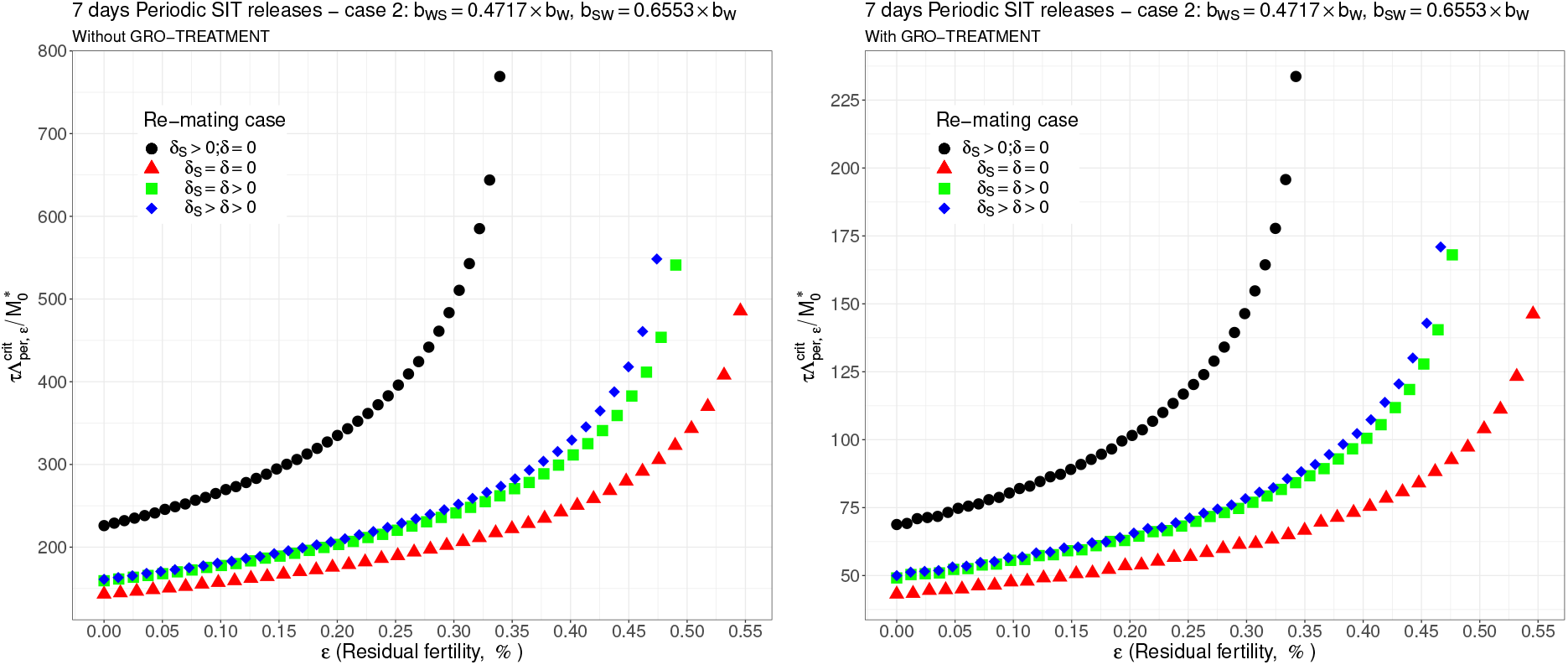
Critical ratio for weekly periodic releases as a function of residual fertility - re-mating case 2 with *b*_*W,S*_ = 0.4717 ×*b*_*W*_ and *b*_*SW*_ = 0.6553 × *b*_*W*_. Simulations with different re-mating configurations: the black bullets, with re-mating rates *δ*_*S*_ > 0 and *δ* = 0; the red triangles, the NO re-mating case, *δ*_*S*_ = *δ* = 0; the green squares, with positive and equal re-mating rates, *δ*_*S*_ = *δ* > 0; the blue diamonds, with positive re-mating rates, *δ*_*S*_ > *δ* > 0.

**Figure 8:**
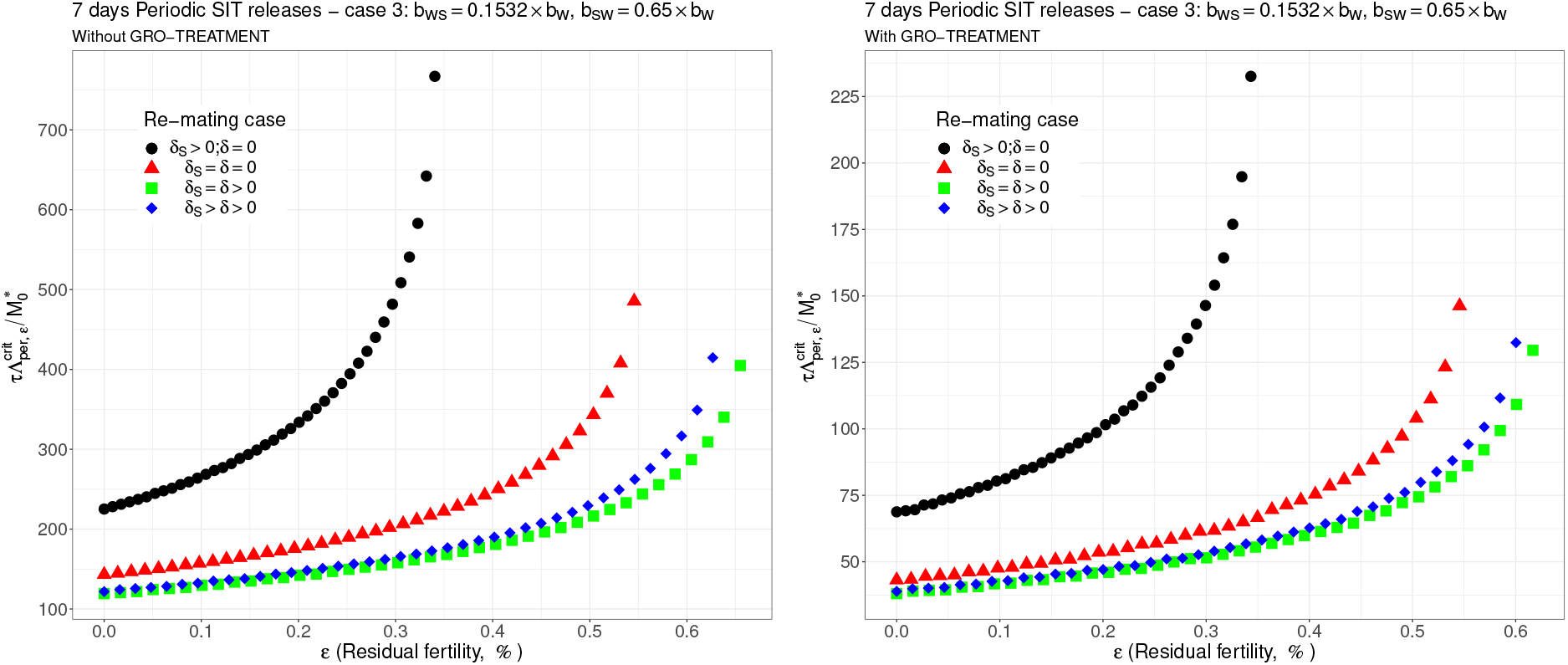
Critical ratio for weekly periodic releases as a function of residual fertility - re-mating case 3 with *b*_*W S*_ = 0.1532 × *b*_*W*_ and *b*_*SW*_ = 0.65 × *b*_*W*_. Simulations with different re-mating configurations: the black bullets, with re-mating rates *δ*_*S*_ > 0 and *δ* = 0; the red triangles, the NO re-mating case, *δ*_*S*_ = *δ* = 0; the green squares, with positive and equal re-mating rates, *δ*_*S*_ = *δ* > 0; the blue diamonds, with positive re-mating rates, *δ*_*S*_ > *δ* > 0.

**Figure 9:**
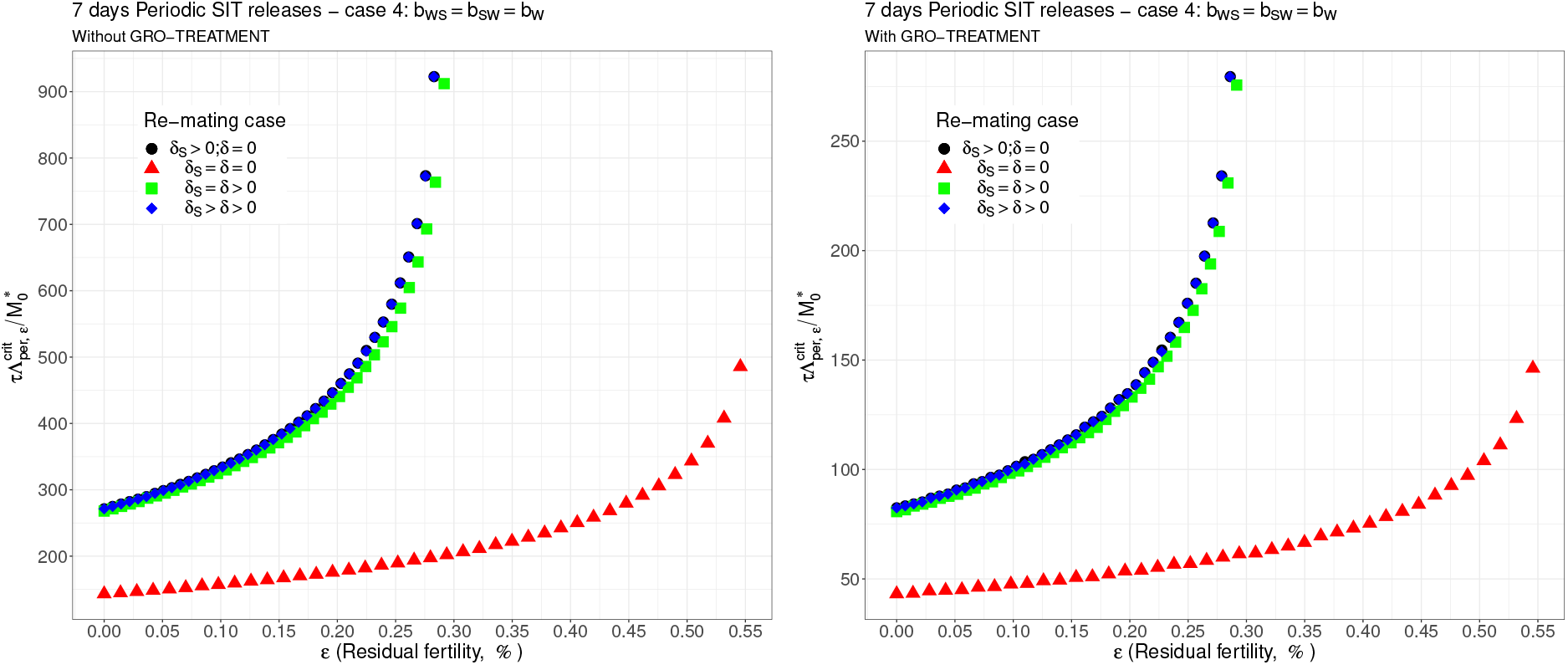
Critical ratio for weekly periodic releases as a function of residual fertility - re-mating case 4 with *b*_*W,S*_ = *b*_*S,W*_ = *b*_*W*_. Simulations with different re-mating configurations: the black bullets, with re-mating rates *δ*_*S*_ > 0 and *δ* = 0; the red triangles, the NO re-mating case, *δ*_*S*_ = *δ* = 0; the green squares, with positive and equal re-mating rates, *δ*_*S*_ = *δ* > 0; the blue diamonds, with positive re-mating rates, *δ*_*S*_ > *δ* > 0.

**Figure 10:**
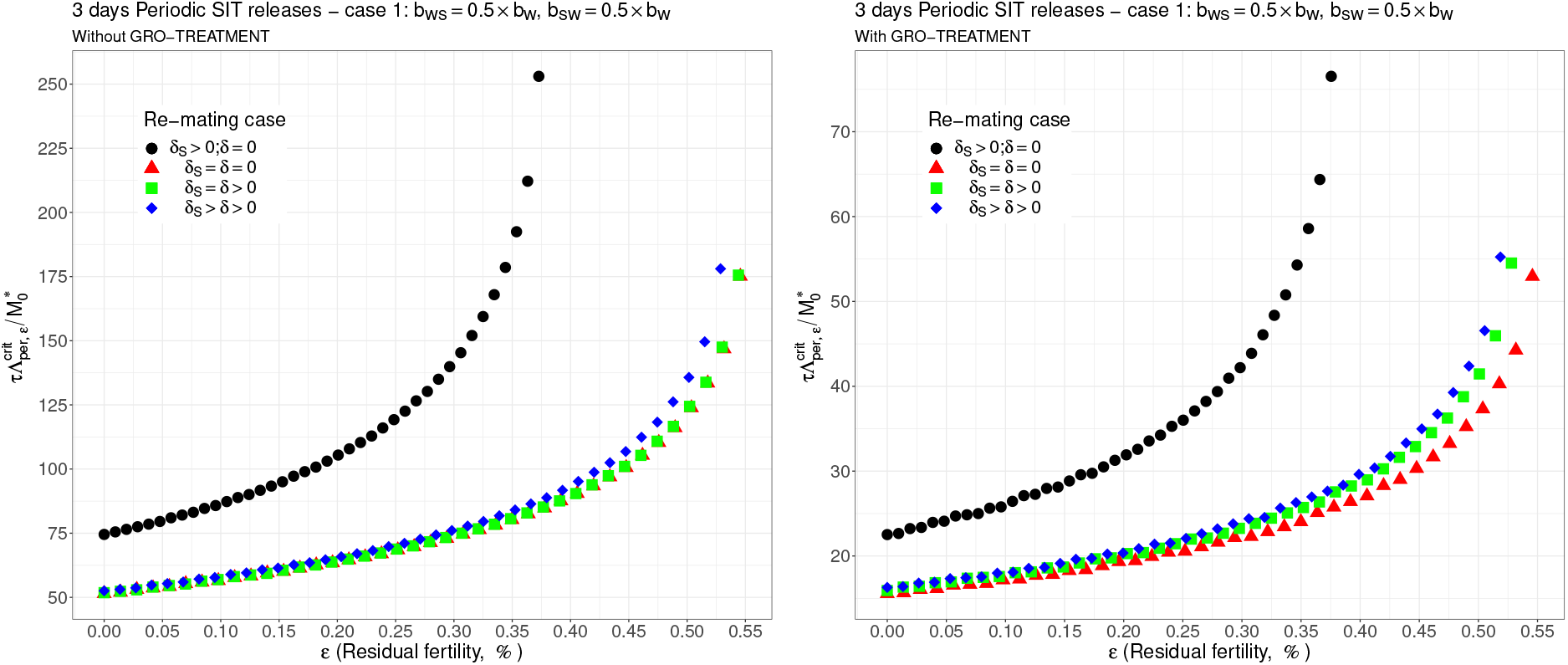
Critical ratio for periodic releases every 3 days based on residual fertility - re-mating case 1 with *b*_*W,S*_ = 0.5 × *b*_*W*_, *b*_*S,W*_ = 0.5 × *b*_*W*_. Simulations with different re-mating configurations: the black bullets, with re-mating rates *δ*_*S*_ > 0 and *δ* = 0; the red triangles, the NO re-mating case, *δ*_*S*_ = *δ* = 0; the green squares, with positive and equal re-mating rates, *δ*_*S*_ = *δ* > 0; the blue diamonds, with positive re-mating rates, *δ*_*S*_ > *δ* > 0.

Last but not least, for periodic releases, whatever the releases period, the gain in sterile males’ releases is only a factor 3 *−* 4 with the GRO treatment compared to without the GRO treatment.

We also derive some simulations to show the dynamics of system (15), page 10, for case 2 with a 3-days SIT release strategy, for different residual fertility, 0% and 0.2%, showing the difference between no-GRO treatment and GRO-treatment: see and compare Figs. 14 and 15 page 24. We choose the size of the release such that 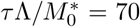: when *ε* = 0, thanks to Figs. 11 page 21, we are above the critical values with and without GRO-treatment. In contrary, when *ε* = 0.002, without GRO-treatment, we are below the critical value, and above when GRO-treatment is used.

**Figure 11:**
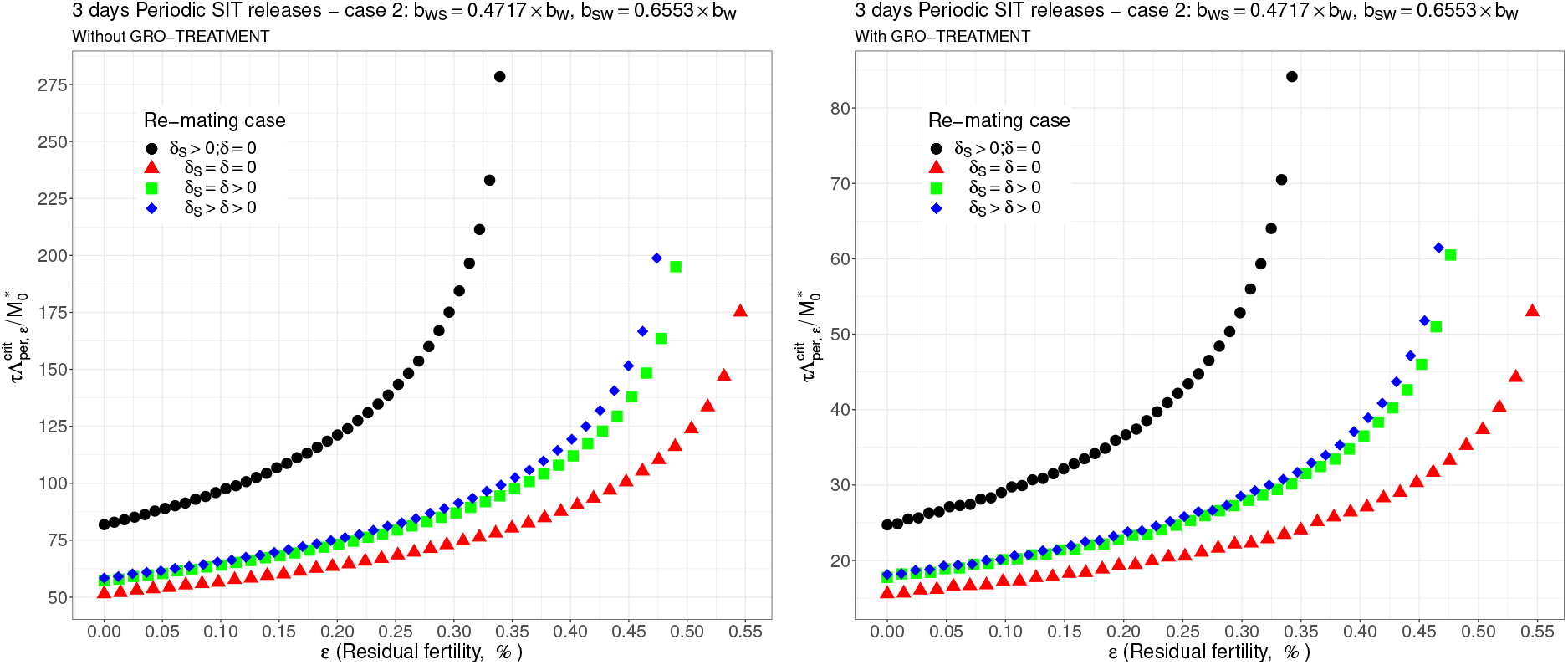
Critical ratio for periodic releases every 3 days based on residual fertility - re-mating case 2 with *b*_*W,S*_ = 0.4717 × *b*_*W*_ and *b*_*SW*_ = 0.6553 × *b*_*W*_. Simulations with different re-mating configurations: the black bullets, with re-mating rates *δ*_*S*_ > 0 and *δ* = 0; the red triangles, the NO re-mating case, *δ*_*S*_ = *δ* = 0; the green squares, with positive and equal re-mating rates, *δ*_*S*_ = *δ* > 0; the blue diamonds, with positive re-mating rates, *δ*_*S*_ > *δ* > 0.

**Figure 12:**
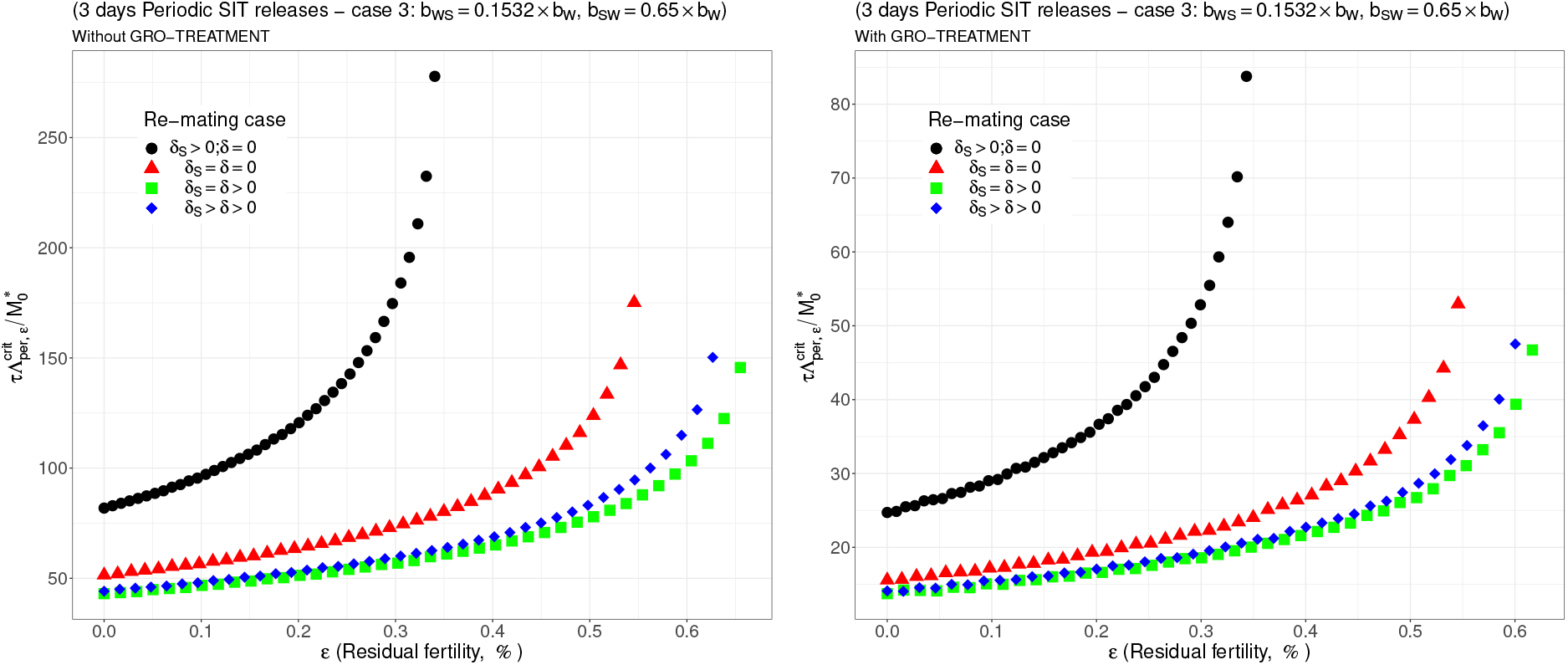
Critical ratio for periodic releases every 3 days based on residual fertility - re-mating case 3 with *b*_*W S*_ = 0.1532 ×*b*_*W*_ and *b*_*SW*_ = 0.65 × *b*_*W*_. Simulations with different re-mating configurations: the black bullets, with re-mating rates *δ*_*S*_ > 0 and *δ* = 0; the red triangles, the NO re-mating case, *δ*_*S*_ = *δ* = 0; the green squares, with positive and equal re-mating rates, *δ*_*S*_ = *δ* > 0; the blue diamonds, with positive re-mating rates, *δ*_*S*_ > *δ* > 0.

**Figure 13:**
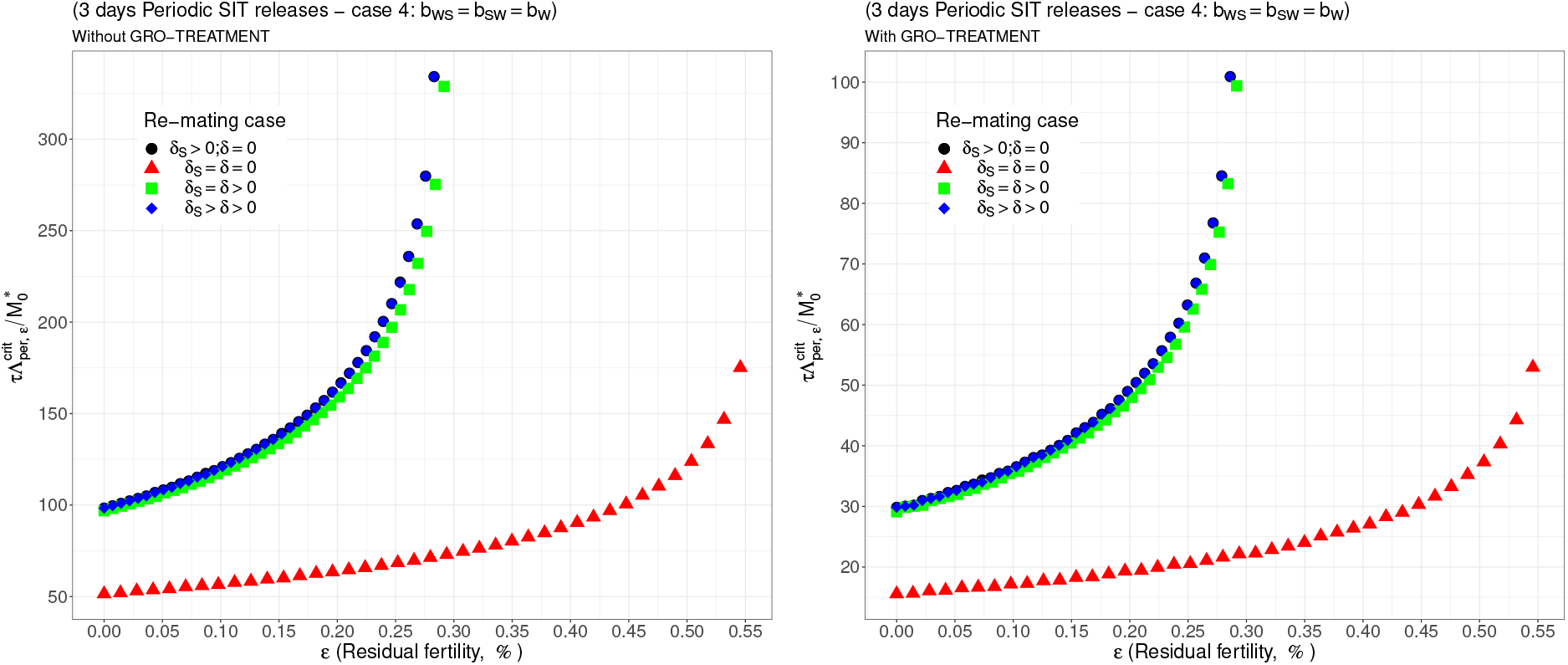
Critical ratio for periodic releases every 3 days based on residual fertility - re-mating case 4 with *b*_*W,S*_ = *b*_*S,W*_ = *b*_*W*_. Simulations with different re-mating configurations: the black bullets, with re-mating rates *δ*_*S*_ > 0 and *δ* = 0; the red triangles, the NO re-mating case, *δ*_*S*_ = *δ* = 0; the green squares, with positive and equal re-mating rates, *δ*_*S*_ = *δ* > 0; the blue diamonds, with positive re-mating rates, *δ*_*S*_ > *δ* > 0.

**Figure 14:**
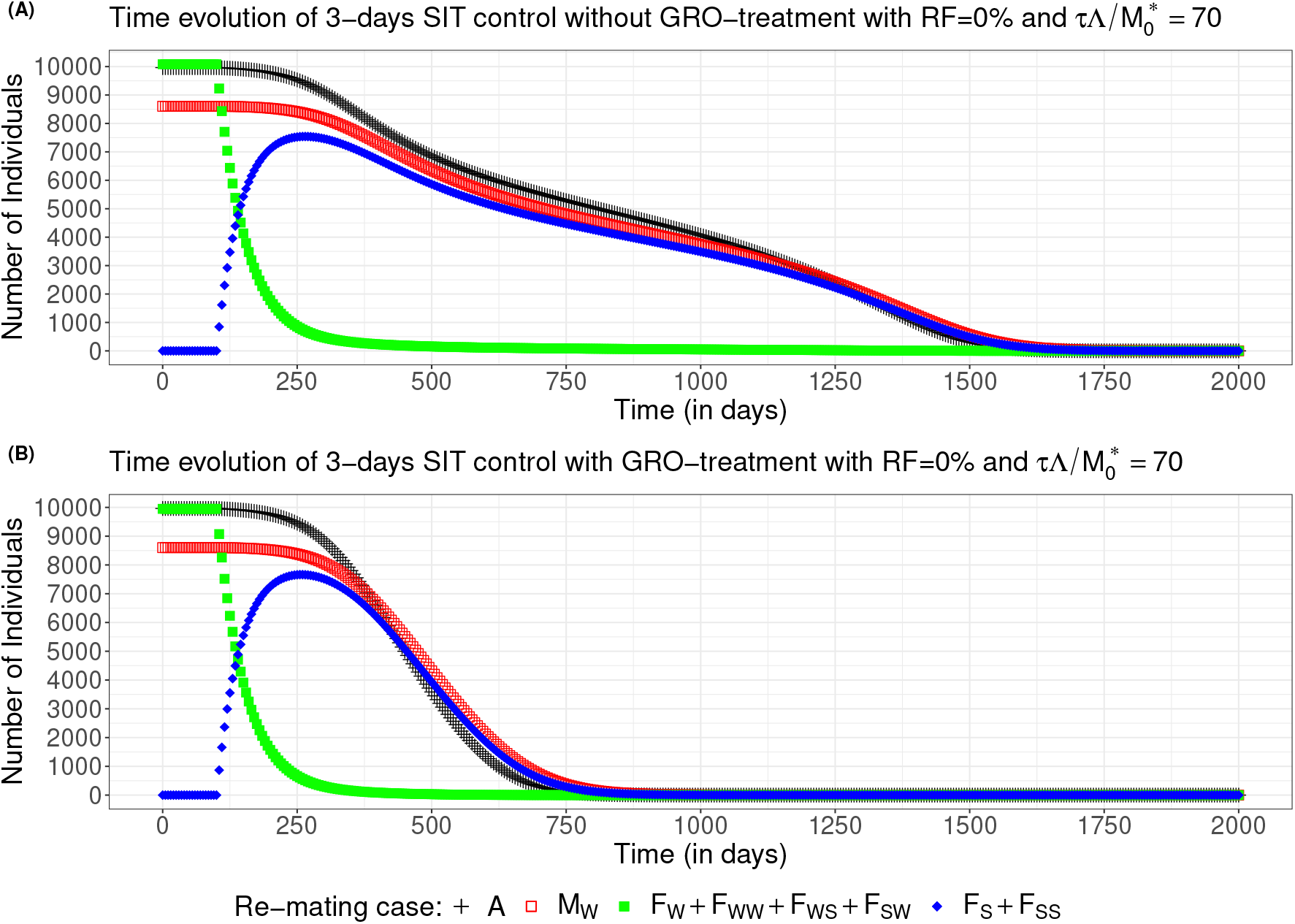
Wild population numbers evolution in time for periodic releases every 3 days and no residual fertility - re-mating case 2 with *b*_*W,S*_ = 0.4717 × *b*_*W*_ and *b*_*SW*_ = 0.6553 ×*b*_*W*_ : (A) Without GRO-treatment (B) With GRO-treatment.

**Figure 15:**
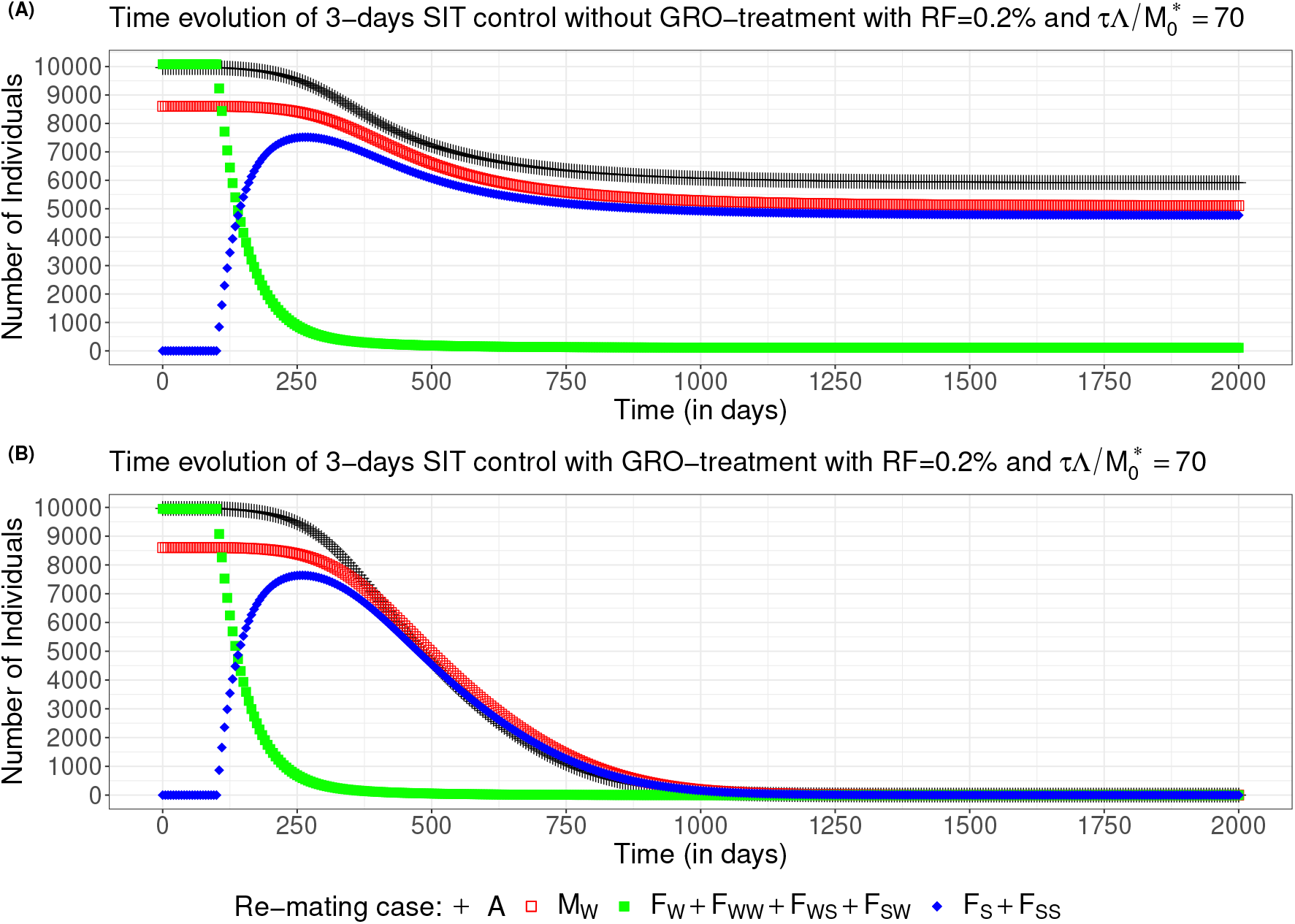
Wild population numbers evolution in time for periodic releases every 3 days and 0.2% residual fertility - re-mating case 2 with *b*_*W,S*_ = 0.4717 ×*b*_*W*_ and *b*_*SW*_ = 0.6553 × *b*_*W*_, : (A) Without GRO-treatment (B) With GRO-treatment.

As expected the dynamics can be rather different: in Fig. 14(A), page 23, the release ratio, 70, is just above the critical release ratio (*≈* 68), that is why it takes so long to reach elimination, while in Fig. 14(B), page 23, elimination is reached in less than 3 years. Even if the wild females (yellow curve) first decay rapidly, during the first 150 days after the starting date, they converge slowly to 0. In the mean time, the sterile females population, *F*_*S*_ + *F*_*SS*_ increase rapidly and then decay following the decay of the wild males population, *M*_*W*_, and the non-adult stages population, *A*. When residual fertility occurs, with *ε* = 0.002, with the same release ratio, we show in Fig. 15(A), page 24, that the system converge to a new positive equilibrium, still with a large population, despite the massive release. When GRO-treatment is used, then elimination is reached in less than 3 years: see Fig. 15(B). It is interesting to note, in Fig. 15(A), that while the amount of fertile females is small (a few hundreds), wild males and non-adult stages are very large. This is due to the fact that the amount of sterile females is large, such that *F*_*S*_ females can potentially, re-mate with wild males and then have offspring. That is why, elimination can only be ensured when the fertile population and also the *F*_*S*_ population have been eliminated.

Of course, once elimination is nearly reached, massive releases are no more necessary: using the strong Allee effect, we can switch to small releases, like in [51], to maintain the wild population very low and to converge slowly but surely to elimination.

Note also that the critical release ratios obtained in the simulations are more or less comparable to the ratios given in [63][Table 3].

## 4 Discussion

The overall success of an SIT program relies on (1) the field efficacy of sterile males in securing matings and sterile progeny and (2) reasonable production costs. The former is affected by several factors: the strain used and mass rearing technique, the radiation dose (and therefore both sterility level and competitiveness of males), the re-mating proportions biased towards wild males. For the latter, we focus on the effect of the release ratio.

Releasing fully sterile males is rarely achieved in SIT programs. Indeed, the dose-response curves for sterility only approach complete sterility asymptotically, and so the elimination of the last 1% residual fertility may require too unreasonably high doses [64]. Thus, the appropriate balance between acceptable residual fertility and quality (that is, competitiveness and average lifespan) must be determined. Our simulations show that the necessary release ratio increases dramatically as residual fertility increases. Moreover, when residual fertility is above a certain threshold, determined by 1*/𝒩*, where 𝒩is the basic offspring number, the SIT may not be an efficient control strategy.

It is sometimes recommended that, at the beginning of SIT deployment, competitive males are preferable to completely sterile males, to ensure that most females mate with the released males. Subsequently, fully sterile males become preferable once the wild population density had decreased to a level where less competitive males would significantly outnumber the wild males, compensating for their lower mating success [23]. Use of this model could help operators better plan their SIT implementation, according to the tolerable limit for residual fertility. More work is needed, however, to understand what is the population reduction target that should trigger the change to releases of fully sterile males.

Competitiveness of sterile males against their wild counterparts is important for the success of SIT programs. The use of GRO treatment to enhance the attractiveness of sterile males is now common in SIT programs against medflies. Our simulations clearly show the positive impact of higher competitiveness values on the success of the release campaign. Even with daily releases, the necessary ratios were almost ten times higher for untreated sterile males.

Our numerical outputs confirm that re-mating has contrasting effects on SIT efficiency, but this mainly depends on the biological parameters, *b*_*_ and *μ*_*_, related to the re-mated females compartments, *F*_*W S*_, *F*_*SW*_ and *F*_*W W*_. For the *F*_*W W*_ compartment, re-mating is beneficial since *b*_*W W*_ > *b*_*W*_ and *μ*_*W W*_ < *μ*_*W*_ [40]. One case is always detrimental to SIT (when *ε*_max_ is low and the release ratio large): it is the case where *δ* = 0 (a wild mated female never re-mate) and *δ*_*S*_ > 0 (a female that mated with a sterile male will re-mate): these are the blue curves in all figures. When re-mating is not beneficial for SIT, then the cases no re-mating and equal re-mating provide the same results, while the case with differential re-mating, *δ*_*S*_ > *δ* > 0, provides smaller values *ε*_max_ a bit, and increases the critical release ratio a bit. However, in the case where re-mating has a positive impact on SIT, then equal re-mating and differential re-mating induce an increase of *ε*_max_ and a decrease of the critical release ratio, compared to the other cases, without re-mating, and with sterile female re-mating only. One case, *b*_*W S*_ = *b*_*SW*_ = *b*_*W*_, seems to be more problematic: this is the case where sterile males have no impact on the oviposition rate as they were not able to transfer sterile sperms, such that the double-mated female only use fertile sperms. Our simulations show that, in that case, SIT control is still possible but only with larger release ratios, and very small residual fertility.

Daily versus periodic releases have a considerable impact on the cost of an operational program. Because of the short average lifespan of sterile males considered in the simulations (see Table 2, page 13), a 7 days periodic releases do not appear suitable. Although lifespan is difficult to estimate in the field, one can consider that sterile males are only useful as long as they can inseminate wild females. However, Abraham et al [25] showed that sperm-depleted sterile males would still be able to transfer accessory gland substances that would trigger a refractory behavior in females.

Our model shows the importance of knowing the impact of re-mating because wild and sterile males may not have the same capacity to inseminate females. Costa et al. [65] showed that a (wild) male would use most of its sperm contained in the testes to inseminate one female and a 24h period appears sufficient to replenish the testes with mature sperm [66]. Wild males can inseminate females every day. On the other hand, a sterile male loses the capacity to mature new sperm as the immature cells would have been damaged by the irradiation process; Catala-Oltra et al. [41] showed that 70% of sterile (irradiated at 100 Gy) males mated again 24h after their first mating, and 25% of males mated up to five times. Sterile Queensland flies were reported to have a maximum of 2 or 3 mating capacity [67]; however, even depleted, they were able to copulate and to induce refractoriness in females. Studies have shown that the injection of male accessory gland substances was able to induce refractoriness in Queensland flies, but also in medflies [25, 68]. Costa et al. [65] showed that a lower quantity of sperm transferred would increase the chances of re-mating in females. A more precise understanding of sterile male reproductive capacity could allow optimization of release frequency, ensuring that mating sterile males always saturates the field. While refractoriness studies are useful, our model, and its analysis, shows that it would be equally important to better know about the biological parameters related to double-mated females, like *b*_*W S*_, *b*_*SW*_, *μ*_*W S*_, and *μ*_*SW*_, and also *b*_*W W*_ and *μ*_*W W*_. As showed in (11), we need to estimate these parameters in order to derive the maximal residual fertility, *ε*_max_, and, thus, the critical release ratio.

## 5 Conclusion

The aim of this study was to investigate the impact of re-mating and residual fertility on sterile insect technique efficiency. Assuming double-mating, in order to ensure existence of a strong Allee effect, i.e. equilibrium 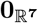 being, at least, locally asymptotically stable, we show that the residual fertility parameter, *ε*, has to be lower than a threshold parameter, *ε*_max_, defined in (11), page 8, that depend on the basic offspring number, 𝒩 (*δ*), and also parameters related to double-mated females. From that point of view, our result also encompasses the result found in [16], when no-remating occurs, where we found that *ε* has to verify *ε𝒩* (0) < 1. Thanks to (11), the larger the basic offspring number, the lower *ε*_max_. However, other parameters are also important in (11): the re-mating rates parameters, *δ* and *δ*_*S*_, and also the parameters *b*_*_ and *μ*_*_ related to the single- and double-mated fertile females, *F*_*W*_, *F*_*W W*_, *F*_*W S*_, and *F*_*SW*_.

Our theoretical and numerical results also show that re-mating is, in general, not beneficial for SIT, in particular when residual fertility occurs. As the level of residual fertility is related to the radiation dose, our results indicate that a small percentage of residual fertility is not necessarily detrimental to SIT. This is important because the radiation dose has an impact on the lifespan and competitiveness of sterile males. When re-mating is not equal, that is, re-mating is faster after mating with a sterile male, it is necessary to drastically increase the release ratiorate of sterile males.

However, we show, in case 3, that re-mating can be beneficial for SIT by increasing *ε*_max_, in particular when *b*_*W S*_ is substantially smaller than *b*_*W*_. Case 3 is based on a very particular case where the sterile male is from a different genotype. This case could occur in ongoing field studies, where the sterile flies, produced far away from the release place, are not necessary of the same genotype than the local fruit flies.

Our results also show that GRO-treatment is very important to enhance the competitiveness of sterile males and allows reducing the release ratios.

To summarize, our study showed that biological parameters related to re-mating, *δ*_*_, *b*_*_, and *μ*_*_ are very important to determine an upper bound for the residual fertility and also the critical release threshold. We found several fragmented information in the literature about these parameters. It seems that it would be more than necessary to estimate all these parameters in order to guarantee or enhance the SIT efficacy. We consider only single and double mating, but our model can be extended to more re-mating, but then, we should be capable to get data related to all re-mating. Not so easy.

This study was performed within the framework of an SIT feasibility project, CeraTIS-Corse in Corsica on a 400 ha area of citrus and stone fruits. The Corsican agricultural context is a mosaic of small plots with different cultivars that span the harvest over several months. The climate is particularly suitable for medflies, which are found in traps from May to December, with low adult levels potentially hibernating and reproducing in pomelos as early as February. Medfly life parameters are known to vary according to the host fruit species, and models can consider this variability according to fruit phenology and temperature variation, when available, over the year to obtain a picture of the natural evolution of the population, as in [43] for *Aedes albopictus* in Réunion island. SIT models consider more simplistic population dynamics with averaged parameters over the year; however, it would be interesting to understand whether variation of released male numbers over the year brings a better or similar control while reducing the program costs. To go further, and in order to optimize the releases ratios, feedback control or mixed-control can be used, like in [21, 16], to adapt the size of the releases along the SIT control, and thus the total cost of the SIT treatment.

Last but not least, it would be interesting to study another SIT control objective, like reducing the wild population below a given economic threshold, like in [43] where decaying the epidemiological risk, i.e. reducing ℛ_0_, the basic reproduction number, below 1, was possible even if *ε 𝒩* > 1, in the case where no re-mating occurs

No operational SIT program exists in France yet; however, there is growing interest in making this tool available for farmers as a response to environmental requirements and to the removal of chemical pesticides. The cost-efficiency of SIT will be a crucial element in growers’ adoption of the technique. Therefore, understanding the optimal release strategy that will reduce production costs while ensuring high field efficacy is key. To date, there is no sterile flies production capacity in France, therefore SIT implementation would rely on import from a producing country. It is therefore crucial to determine the limiting factors that may affect release success. This study has shown that field operators should try to gather a better understanding of re-mating occurrence in the field, but should also carefully specify or control the accuracy of the sterilizing dose and level of residual fertility of the imported sterile flies. Our model being generic, it can be applied to other fruit flies or pests, and, if necessary, with more re-mating.

## Acknowledgments

First of all, we would like to deeply thank the reviewers and the handling editor for their thorough reviews, that greatly improved the initial manuscript.

The authors acknowledge the financial support of the CeraTIS - Corse project, within the framework of ECOPHYTO 2019, “Leviers territoriaux pour réduire l’utilisation et les risques liés aux produits phytopharmaceutiques (2020 - 2023)”. YD is also co-funded by the European Union: Agricultural Fund for Rural Development (EAFRD), the European Regional Development Fund (ERDF),by the Departmental council of La Réunion and by the Regional council of la Réunion. YD acknowledges the support of the DST/NRF SARChI Chair, South Africa M3B2 in Mathematical Models and Methods in Biosciences and Bioengineering at the University of Pretoria, South Africa (grant 82770). YD also acknowledges the support of the STIC AmSud project BIO-CIVIP 23-STIC-02.

## A Appendix A Proof of Theorem 1

We compute the Jacobian of system (4)

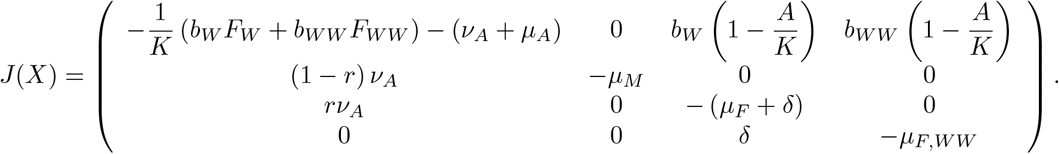

Then we set

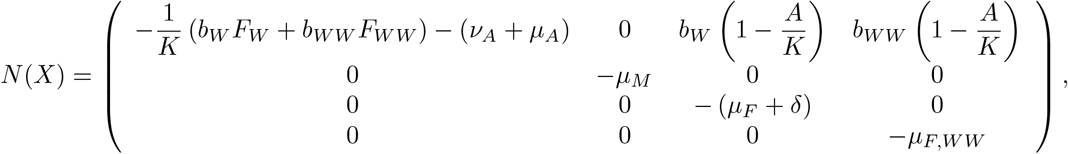

and

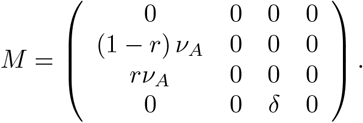

Matrix *N* is Metzler stable and *M* a nonnegative matrix, such that *M* + *N* is a regular splitting. Then

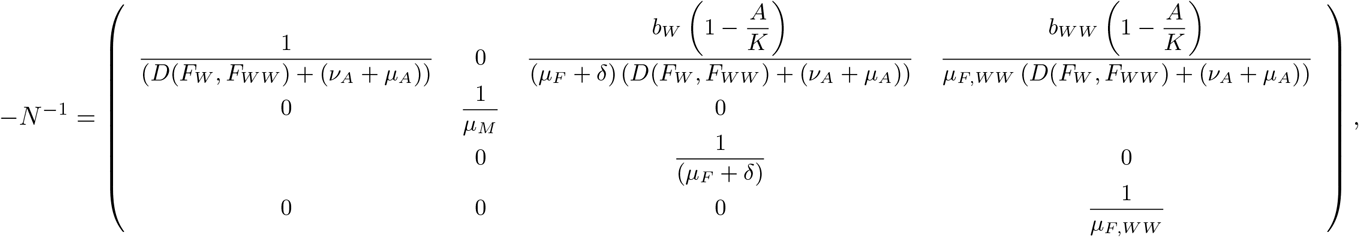

with 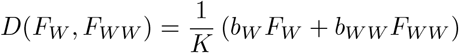. Thus

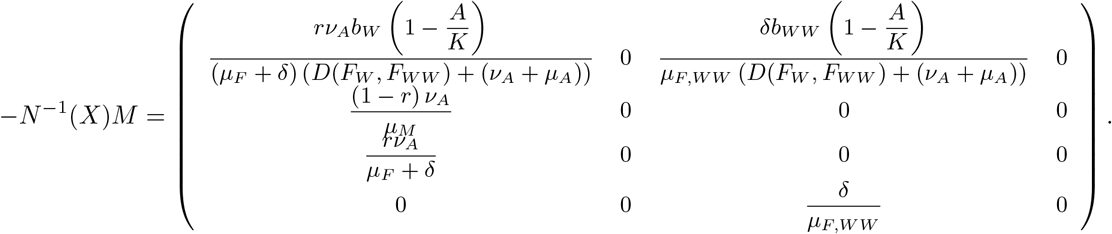

Thus, the characteristic polynomial becomes

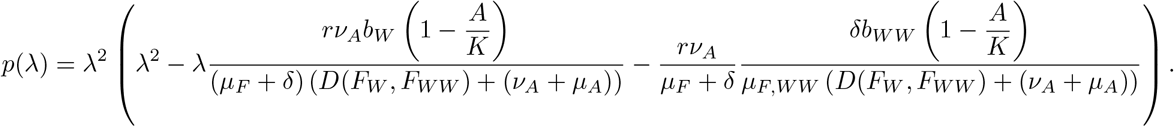

We recall that, for a second order polynomial *q*(*z*) = *z*^2^ + *a*_1_*z* + *a*_2_, to show that all roots are in the inside the unit disc, the Jury criterion leads to the following necessary conditions

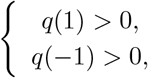

and a sufficient condition

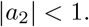

- When *E* = **0**, then

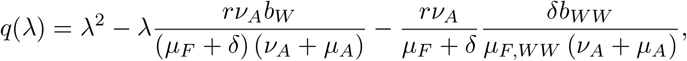

such that

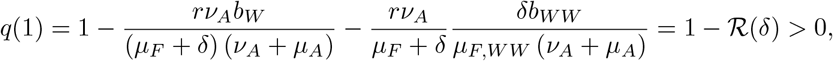

when *ℛ* (*δ*) < 1. From *q*(1) > 0, we deduce that

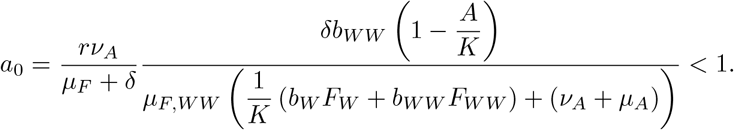

In addition

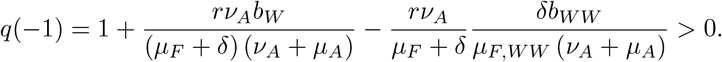

Thus, we deduce that *ρ*(*−N* ^*−*1^(**0**)*M*) < 1, which implies that *s*(*J* (**0**) < 0. Thus the trivial equilibrium is **0** is LAS iff *ℛ* (*δ*) < 1 for system (4).
- Assuming ℛ> 1, then the endemic equilibrium **E** exists. Using the equilibrium values given in (7)-(8)-(9)-(10), page 8, the characteristic equation becomes

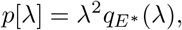

with

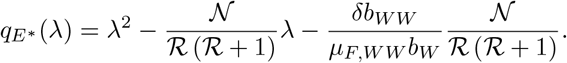

We have

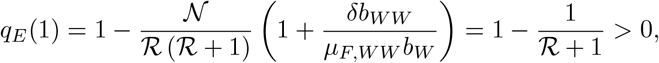

because *ℛ* > 0, which implies that

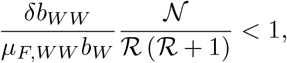

and also

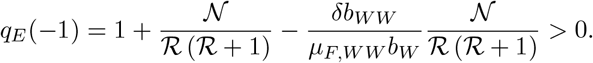

Thus, we deduce that *ρ*(*−N* ^*−*1^(**E**^*^)*M*) < 1, which implies that *s*(*J* (**E**^*^) < 0. Thus the trivial equilibrium is **E**^*^ is LAS iff *ℛ* (*δ*) > 1 for system (4).

From the Theory of Monotone Cooperative system [49], and following [69, Theorem 6] or [51, Theorem 1], we know that once an equilibrium exists, is unique and LAS, then it is GAS. Thus, our results follow.

## B Appendix B SIT model - Stability analysis of Equilibrium 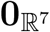

We compute the Jacobian related to system (2) at 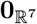

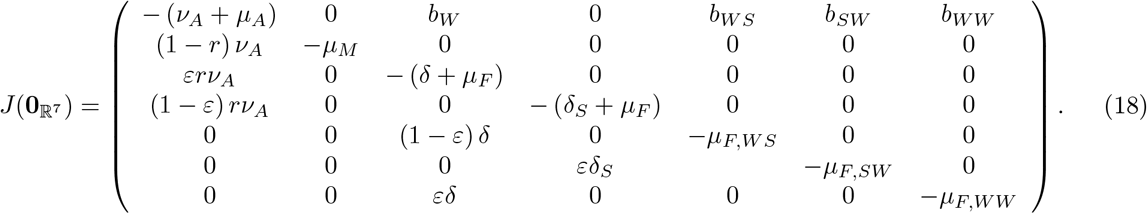

Setting

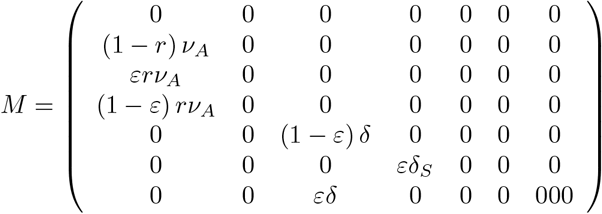

and

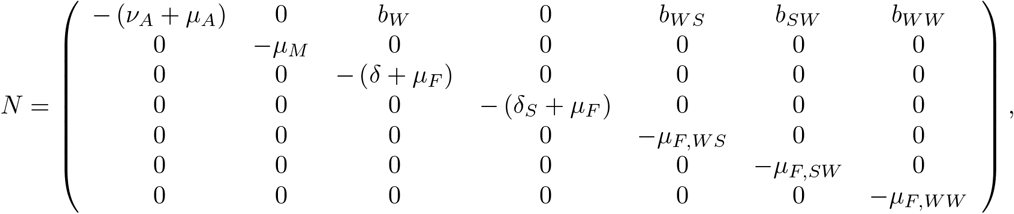

where *N* is an invertible M-matrix and *M* a nonnegative matrix, then, we deduce

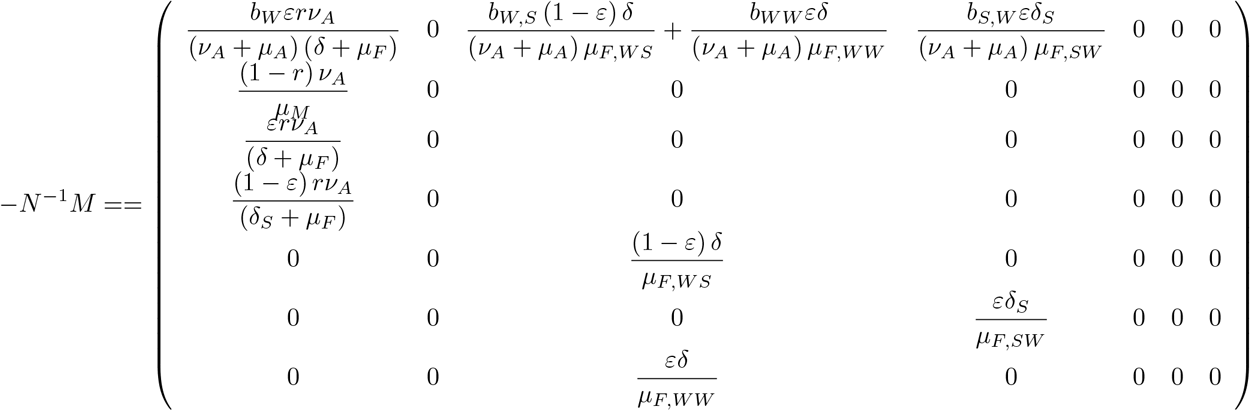

After some computations, we can show that the characteristic polynomial of *−N* ^*−*1^*M* reduces to *− λ*^5^*q*(*λ*), where

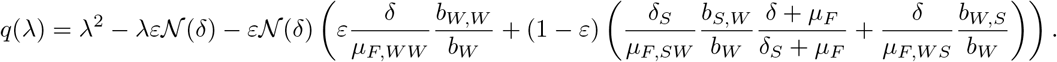

To show that all roots, *λ*^*^, are such that *Iλ*^*^*I* < 1, we have to check Jury’s criterion (recalled in Appendix A, page 30), namely *q*(1) > 0, *q*(*−*1) > 0 and

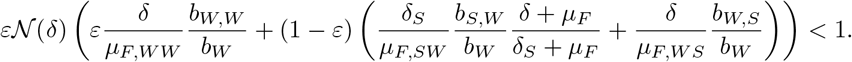

It is easy to check that *q*(*−*1) > 0. Then

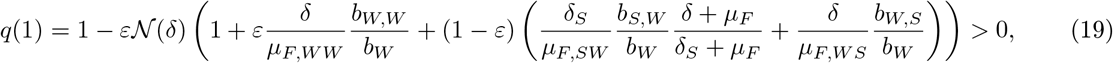

iff

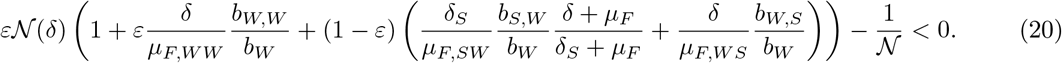

Thus we have to find under what condition the previous polynomial in *ε* takes negative values. After straightforward calculations, we found that for 0 *≤ ε ≤ ε*^*^, such that

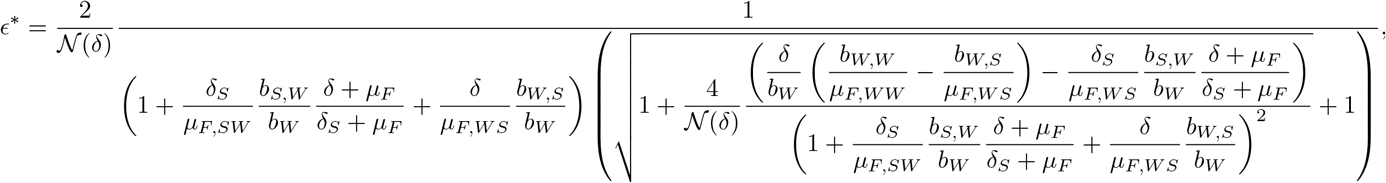

condition (20) is verified. Thus we conclude that *ρ*(*− N* ^*−*1^*M*) < 1 which implies 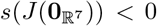, and the result follows.

## C Appendix C SIT model - looking for equilibria

We are looking for a condition to have at least one positive equilibrium. Thus, we have to solve

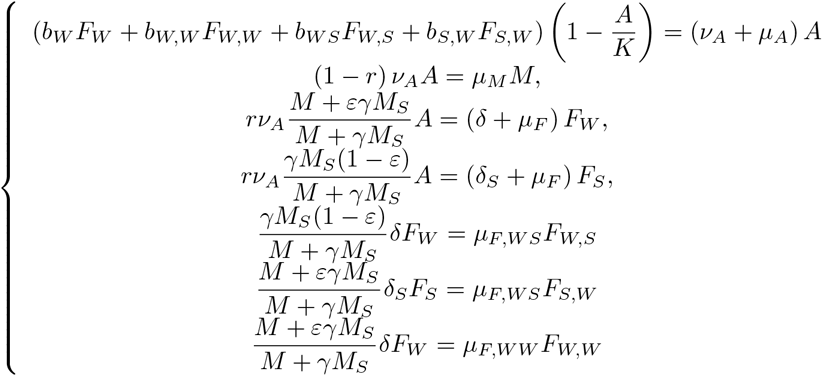

such that

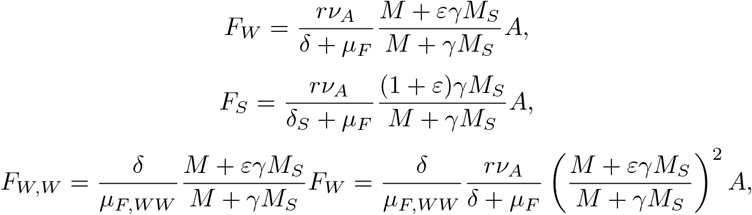

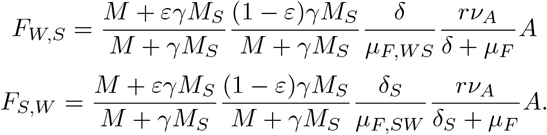

Thus from the first equation

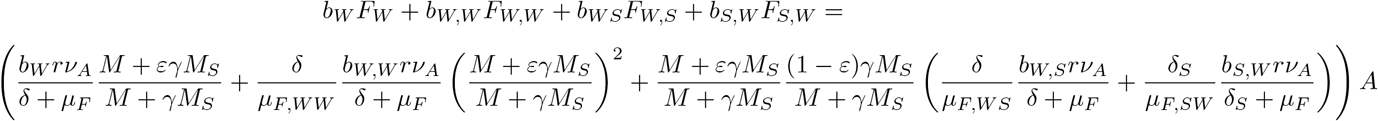

that is

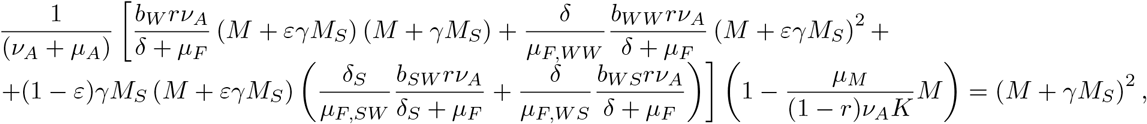

that is

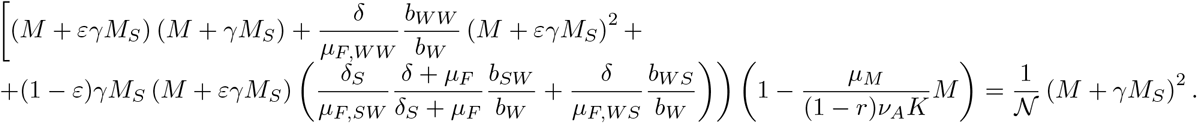

Expanding the left-hand side leads to

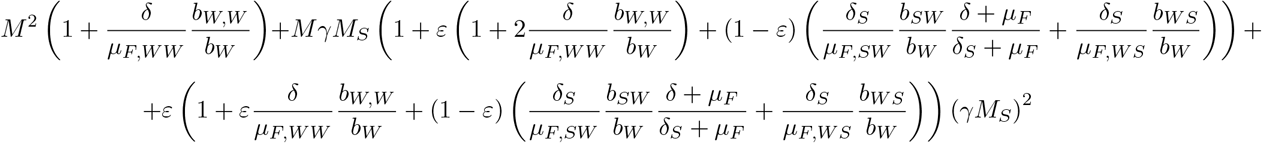

We have

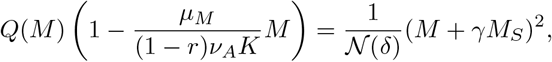

with

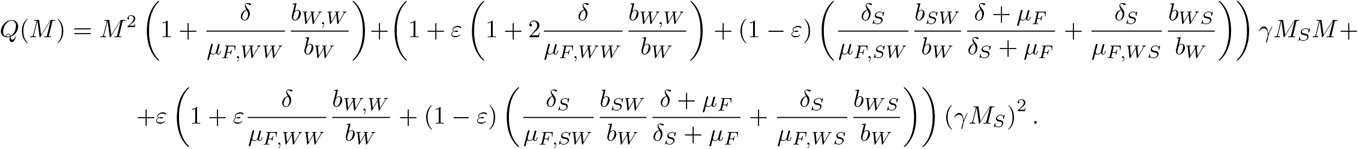

The roots of *Q* are

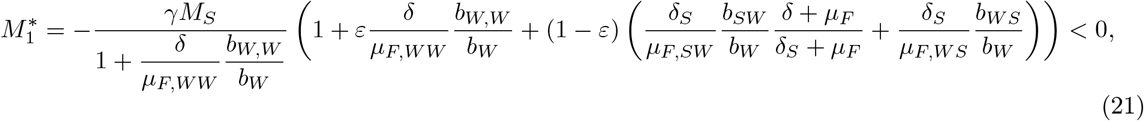

and 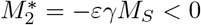. They are both real negative. Thus we are looking for positive roots of

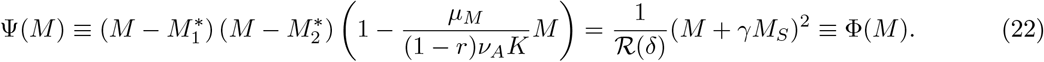

Thus, we can derive 3 cases as showed in Fig. 16, page 35:

**Figure 16:**
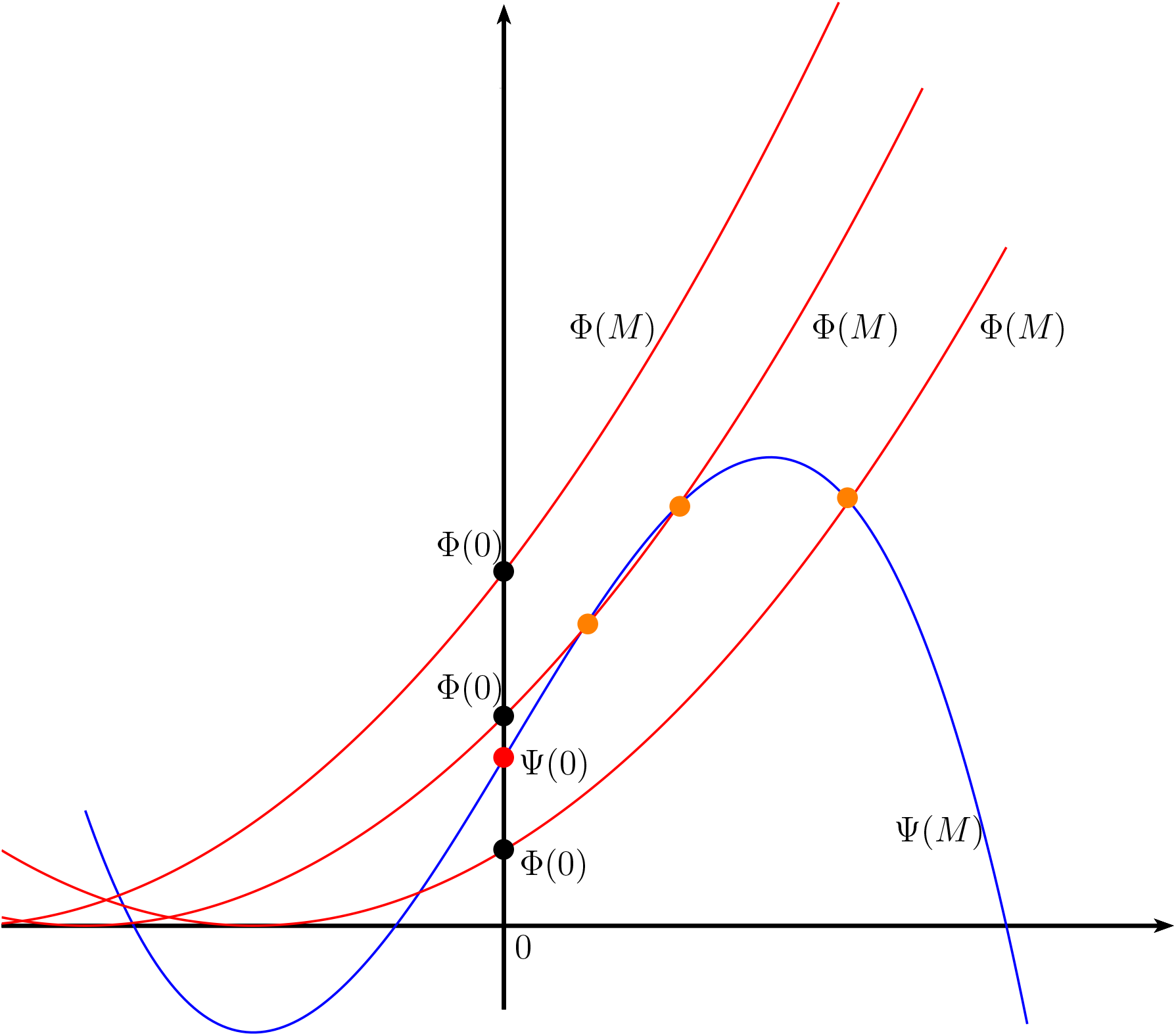
Intersections between Ψ (in blue) and Φ (in red) - 3 cases with no, one or two intersections (orange bullets)

1. Since Φ(0) < Ψ(0), then there always exists a positive root, *M* ^*^. Note also that Φ(0) *≤* Ψ(0) is equivalent to choose *ε* such that

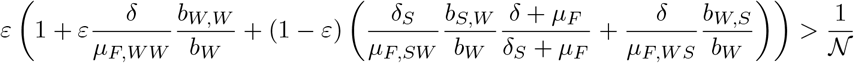

which is the opposite condition of (20), page 33, that is *ε* > *ε*_max_, where *ε*_max_ given in (11), page 8, and for which equilibrium 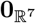 is unstable.
2. Since Φ(0) *≥* Ψ(0), that is *ε* ∈ [0, *ε*_max_], we have two cases
  a. either *γM*_*S*_ is small such that we can have one intersection or two intersections between Ψ and Φ
  b. either *γM*_*S*_ is sufficiently large such that Ψ and Φ do not intersect.

Thus, we can deduce that there exists a critical threshold 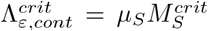 such that above this critical threshold no positive equilibrium can exists, only the trivial equilibrium, i.e. 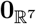. We are not able to derive a formula for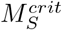, and thus for 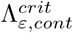, but we can solve (22), page 34, to find it.

In the case where *ε* = *ε*_max_, we can derive an explicit formula for 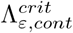. Indeed, we have 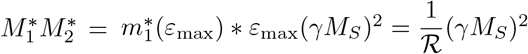, that is 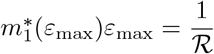 for all *M*_*S*_ > 0, with

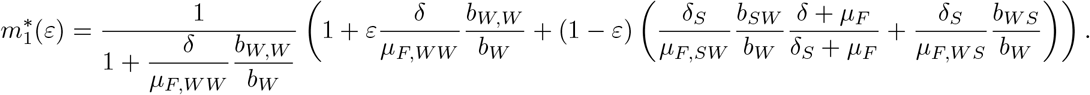

Then, equation (22) becomes

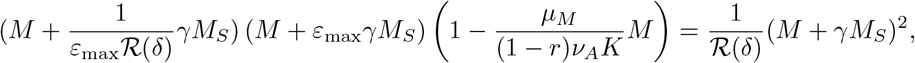

and it reduces to the following, equivalent, second order equation to solve

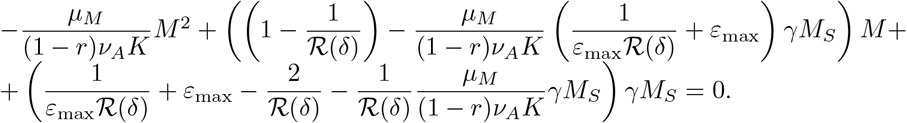

Computing the discriminant leads to

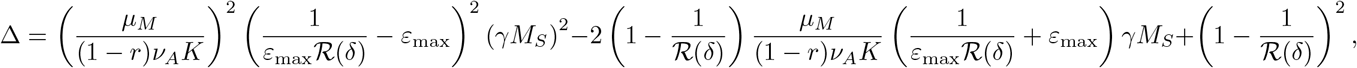

that is a second-order polynomial in *γM*_*S*_, for which the discriminant is

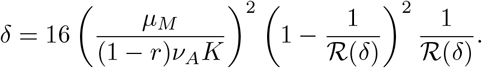

Thus we derive the first positive root

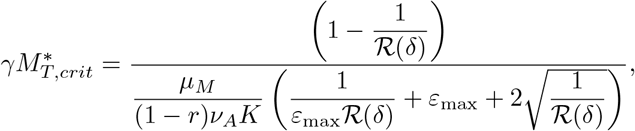

from which we deduce

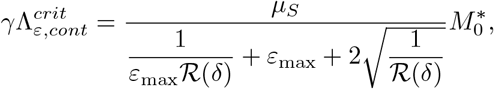

related to the initial male population at equilibrium, 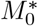.

## D Appendix D Proof of Theorem 2

We first check the condition on *ε* to have 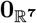 LAS for the monotone cooperative system (3), page 7.

We compute

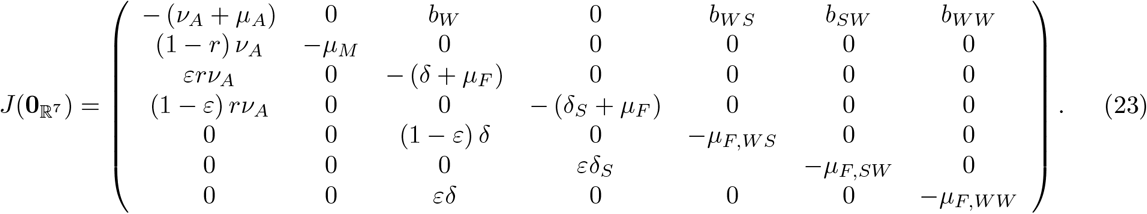

We observe that system (2) and system (3) have the same Jacobian at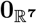, such that, following Lemma 2, page 8, we can deduce that 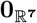 is also LAS for system (3) when *ε ≤ ε*_max_, and unstable otherwise. Following the same reasoning than in appendix C, we show that a positive equilibrium exists for model (3) if there exists positive roots of

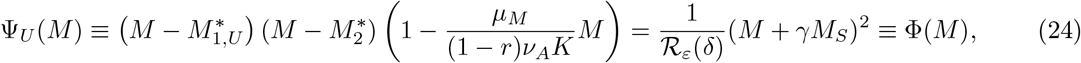

where 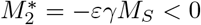 and

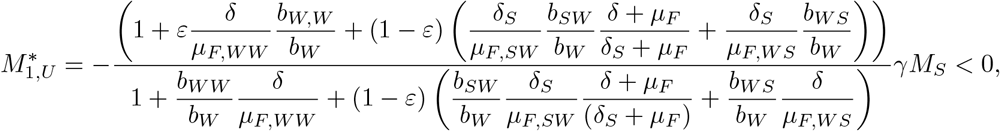

and

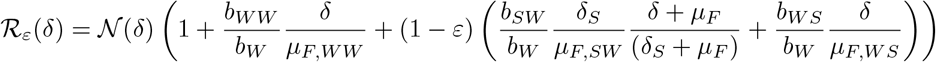

Notice that 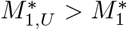, where 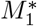 is defined in (21). Like in Appendix C, we derive 3 cases, as showed in Fig. 16, page 35, where Ψ is replaced by Ψ_*U*_.

1. Since Φ_0_(0) < Ψ(0), then (24) always admits a positive root, *M* ^*^. Note also that Φ_*U*_ (0) *≤* Ψ(0) is equivalent to choose *ε* such that

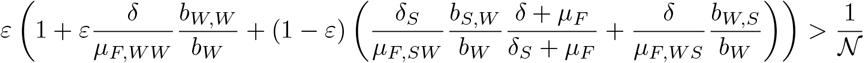

which is the opposite condition of (20), page 33, that is *ε* > *ε*_max_, where *ε*_max_ given in (11), page 8, and for which equilibrium 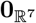 is unstable for system (3).
2. Since Φ_*U*_ (0) *≥* Ψ(0), that is *ε* ∈ [0, *ε*_max_], we have two cases
  a. either *γM*_*S*_ is small such that we can have one intersection or two intersections between Ψ_*U*_ and Φ
  b. either *γM*_*S*_ is sufficiently large such that Ψ_*U*_ and Φ do not intersect.

Thus, we can deduce that there exists a critical threshold 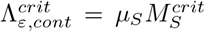 such that above this critical threshold no positive equilibrium can exists, only the trivial equilibrium, i.e. 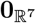. We are not able to derive a formula for 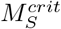, and thus for 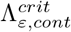, but we can solve (24), page 36, to find it.

Assume *ε* ∈ [0, *ε*_max_], and Λ, the release rate, greater than 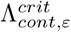, such that according to the previous reasoning, only the trivial equilibrium exists, 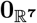, that is also LAS. Thanks to the Theory of Monotone Cooperative system [49], and following [69, Theorem 6] or [51, Theorem 1], it is straightforward to deduce that 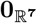 is not only LAS but it is GAS when *ε ≤ ε*_max_. The equilibrium 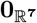 being GAS for the auxiliary Monotone system (3), it is also GAS for system (2), for 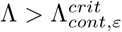.

## References

[1] V. A. Dyck, J. Hendrichs, and A. S. Robinson. The Sterile Insect Technique, Principles and Practice in Area-Wide Integrated Pest Management. Springer, Dordrecht, 2006.

[2] E. F. Knipling. Possibilities of insect control or eradication through the use of sexually sterile males. Journal of Economic Entomology, 48(4):459–462, 1955.

[3] A.S. Robinson. Genetic sexing strains in medfly, ceratitis capitata, sterile insect technique programmes. Genetica, 116:5–13, 2002.

[4] C. Cáceres, K. Bourtzis, G. Gouvi, Vreysen M.J.B., N.S.B. Somda, M. Hejničková, F. Marec, and Meza J.S. Development of a novel genetic sexing strain of ceratitis capitata based on an x-autosome translocation. Sci. Rep., 13, 2023.

[5] I. Plá, J. Garcia de Oteyza, C. Tur, M.A. Martinez, M.C. Laurin, E. Alonso, M. Martinez, A. Martin, R. Sanchis, M.C. Navarro, M.T. Navarro, R. Argilés, M. Briasco, Ó. Dembilio, and V. Dalmau. Sterile insect technique programme against mediterranean fruit fly in the valencian community (spain). Insects, 12(5), 2021.

[6] R. Asadi, R. Elaini, R. Lacroix, T. Ant, A. Collado, L. Finnegan, P. Siciliano, A. Mazih, and M. Koukidou. Preventative releases of self-limiting ceratitis capitata provide pest suppression and protect fruit quality in outdoor netted cages. International Journal of Pest Management, 66(2):182–193, 2020.

[7] F. Landmann. The wolbachia endosymbionts. Microbiol Spectr., 7(2), 2019.

[8] S. Zabalou, M. Riegler, M. Theodorakopoulou, C. Stauffer, C. Savakis, and K. Bourtzis. Wolbachia-induced cytoplasmic incompatibility as a means for insect pest population control. Proc. Natl. Acad. Sci., 101:15042–15045, 2004.

[9] T. Shelly and D. McInnis. Sterile Insect Technique and Control of Tephritid Fruit Flies: Do Species With Complex Courtship Require Higher Overflooding Ratios? Annals of the Entomological Society of America, 109(1):1–11, 2015.

[10] W.R. Enkerlin. Impact of fruit fly control programmes using the sterile insect technique. In V A Dyck, Jorge Hendrichs,, and A. S. Robinson, editors, Sterile Insect Technique. Principles and Practice in Area-Wide Integrated Pest Management, pages 651–676. CRC Press, 2005.

[11] P. Van den Driessche. Some analytical models for biotechnical methods of pest control. In V.A. Dyck, J. Hendrichs, and A.S. Robinson, editors, Pest Control: Operations and Systems Analysis in Fruit Fly Management. Springer Berlin Heidelberg, 1985.

[12] J. Li and Z. Yuan. Modelling releases of sterile mosquitoes with different strategies. Journal of Biological Dynamics, 9(1):1–14, 2015.

[13] H.J. Barclay. Mathematical models for using sterile insects. In V.A. Dyck, J. Hendrichs, and A.S. Robinson, editors, Sterile Insect Technique: Principles And Practice In Area-Wide Integrated Pest Management (2nd ed.). CRC Press, 2021.

[14] M. Strugarek, H. Bossin, and Y. Dumont. On the use of the sterile insect technique or the incompatible insect technique to reduce or eliminate mosquito populations. Applied Mathematical Modelling, 68:443–470, 2019.

[15] M. Huang, X. Song, and J. Li. Modelling and analysis of impulsive releases of sterile mosquitoes. Journal of Biological Dynamics, 11(1):147–171, 2017.

[16] MS Aronna and Y Dumont. On nonlinear pest/vector control via the sterile insect technique: Impact of residual fertility. Bulletin of Mathematical Biology, 82(8):110, Aug 2020.

[17] L. Esteva and H. Mo Yang. Mathematical model to assess the control of aedes aegypti mosquitoes by the sterile insect technique. Mathematical Biosciences, 198(2):132–147, 2005.

[18] R. Anguelov, Y. Dumont, and J. Lubuma. On nonstandard finite difference schemes in biosciences. AIP Conf. Proc., 1487:212–223, 2012.

[19] Y. Dumont and J. M. Tchuenche. mathematical studies on the sterile insect technique for the chikungunya disease and aedes albopictus. Journal of Mathematical Biology, 65 (5):809–854, 2012.

[20] R.C.A Thomé, M. Mo Yang, and L. Esteva. Optimal control of aedes aegypti mosquitoes by the sterile insect technique and insecticide. Mathematical Biosciences, 223(1):12–23, 2010.

[21] P.-A. Bliman, D. Cardona-Salgado, Y. Dumont, and O. Vasilieva. Implementation of control strategies for sterile insect techniques. Mathematical Biosciences, 314:43–60, 2019.

[22] L. Almeida, M. Duprez, Y. Privat, and N. Vauchelet. Optimal control strategies for the sterile mosquitoes technique. Journal of Differential Equations, 311:229–266, 2022.

[23] A. Parker and K. Mehta. Sterile insect technique: a model for dose optimization for improved sterile insect quality. Florida Entomologist, 90(1):88–95, 2007.

[24] FAO/IAEA/USDA. Product Quality Control for Sterile Mass-Reared and Released Tephritid Fruit Flies. FAO/IAEA Programme of Nuclear Techniques in Food and Agriculture. IAEA, Vienna, Austria, 2019.

[25] S. Abraham, V. Diaz, A. Moyano, G. Castillo, J. Rull, L. Suárez, A.F. Murúa, V. Pantano, D. Molina, and S.M. Ovruski. Irradiation dose does not affect male reproductive organ size, sperm storage, and female remating propensity in Ceratitis capitata. Bulletin of Entomological Research, 111(1):82–90, 2021.

[26] A. S. Robinson, J. P. Cayol, and J. Hendrichs. Recent findings on medfly sexual behavior: implications for SIT. Florida Entomologist, 85(1):171–181, 04 2002.

[27] S. A. Lux, J. C. Vilardi, P. Liedo, K. Gaggl, G. E. Calcagno, F. N. Munyiri, M. T. Vera, and F. Manso. Effects of irradiation on the courtship behavior of medfly (Diptera, Tephritidae) mass reared for the sterile insect technique. Florida Entomologist, 85(1):102–112, 2002.

[28] M. Bonizzoni, L.M. Gomulski, S. Mossinson, C.R. Guglielmino, A.R. Malacrida, B. Yuval, and G. Gasperi. Is Polyandry a Common Event Among Wild Populations of the Pest Ceratitis capitata? Journal of Economic Entomology, 99(4):1420–1429, 2006.

[29] M. Bonizzoni, B.I. Katsoyannos, R. Marguerie, C.R. Guglielmino, G. Gasperi, A. Malacrida, and T. Chapman. Microsatellite analysis reveals remating by wild Mediterranean fruit fly females, Ceratitis capitata. Molecular Ecology, 11(10):1915–1921, 2002-10.

[30] R. Morelli, B.J. Paranhos, A. M. Coelho, R. Castro, L. Garziera, F. Lopes, and J.M.S. Bento. Exposure of sterile mediterranean fruit fly (diptera: Tephritidae) males to ginger root oil reduces female remating. Journal of Applied Entomology, 137(1):75–82, 2013.

[31] F.W. Avila, L.K. Sirot, B.A. LaFlamme, C.D. Rubinstein, and M.F. Wolfner. Insect seminal fluid proteins: identification and function. Annual Review of Entomology, 56:21–40, 2011.

[32] C. Gillott. Male accessory gland secretions: modulators of female reproductive physiology and behavior. Annual Review of Entomology, 48(1):163–184, 2003.

[33] Takahisa Miyatake, Tracey Chapman, and Linda Partridge. Mating-induced inhibition of remating in female mediterranean fruit flies ceratitis capitata. Journal of Insect Physiology, 45(11):1021–1028, 1999.

[34] M.T. Vera, J.L. Cladera, G. Calcagno, J.C. Vilardi, and D.O. McInnis. Remating of Wild Ceratitis capitata (Diptera: Tephritidae) Females in Field Cages. Annals of the Entomological Society of America, 96(4):563–570, 07 2003.

[35] D. Marchini, G. del Bene, L.F. Falso, and R. Dallai. Structural organization of the copulation site in the medfly Ceratitis capitata (Diptera: Tephritidae) and observations on sperm transfer and storage. Arthropod Structure & Development, 30(1):39–54, 2001-10.

[36] S.G. Lee, S.D. McCombs, and S.H. Saul. Sperm precedence of irradiated mediterranean fruit fly males (diptera: Tephritidae). Proceedings of the Hawaiian Entomological Society, 36:47–59, 2003.

[37] K. P. Katiyar and E. Ramirez. Mating Frequency and Fertility of Mediterranean Fruit Fly Females Alternately Mated with Normal and Irradiated Males12. Journal of Economic Entomology, 63(4):1247–1250, 08 1970.

[38] Stephen H. Saul and Susan D. McCombs. Dynamics of Sperm Use in the Mediterranean Fruit Fly (Diptera: Tephritidae): Reproductive Fitness of Multiple-Mated Females and Sequentially Mated Males. Annals of the Entomological Society of America, 86(2):198–202, 03 1993.

[39] F. Scolari, B. Yuval, and L.M. et al. Gomulski. Polyandry in the medfly shifts in paternity mediated by sperm stratification and mixing. BMC Genet, 15 (Suppl 2)(S10), 2014.

[40] T. S. Whittier and T. E. Shelly. Productivity of singly vs. multiply mated female mediterranean fruit flies, ceratitis capitata (diptera: Tephritidae). Bulletin of the Entomological Society of America, 66, 1993.

[41] M. Catalá-Oltra, E. Llácer, O. Dembilio, I. Pla, A. Urbaneja, and M. Pérez-Hedo. Remating in Ceratitis capitata sterile males: Implications in sterile insect technique programmes. Journal of Applied Entomology, 145(10):958–965, 2021.

[42] Y. Dumont and I.V. Yatat-Djeumen. Sterile insect technique with accidental releases of sterile females. impact on mosquito-borne diseases control when viruses are circulating. Mathematical Biosciences, 343:108724, 2022.

[43] Y. Dumont and M. Duprez. Modeling the impact of rainfall and temperature on sterile insect control strategies in a tropical environment. Journal of Biological Systems, page 2450012, 2024.

[44] M. Ghabbari and J.M. Ben Jemâa. Influence of different hosts on biological parameters of the fruit fly Ceratitis capitata (Diptera: Tephritidae) in Tunisia. Journal of Oasis Agriculture and Sustainable development, 4(1):48–57, 2022.

[45] W Pieterse, A Manrakhan, J.S. Terblanche, and P. Addison. Comparative demography of bactrocera dorsalis (hendel) and ceratitis capitata (wiedemann) (diptera: Tephritidae) on deciduous fruit. Bulletin of Entomological Research, 110(2):185–194, 2020.

[46] M. Bonizzoni, B.I. Katsoyannos, R. Marguerie, C.R. Guglielmino, G. Gasperi, A. Malacrida, and T. Chapman. Microsatellite analysis reveals remating by wild mediterranean fruit fly females, ceratitis capitata. Molecular Ecology, 11(10):1915–1921, 2002.

[47] K. Kraaijeveld and T. Chapman. Effects of male sterility on female remating in the mediterranean fruitfly, ceratitis capitata. Proc Biol Sci., 271(Suppl 4), 2004.

[48] B. J. Paranhos, D. McInnis, R. Morelli, R. M. Castro, L. Garziera, L. G. Paranhos, K. Costa, C. Gava, M. L. Z. Costa, and J. M. M. Walder. Optimum dose of ginger root oil to treat sterile mediterranean fruit fly males (diptera: Tephritidae). Journal of Applied Entomology, 137(1):83–90, 2013.

[49] H. Smith. Monotone dynamical systems: An introduction to the theory of competitive and cooperative systems. American Mathematical Society, Providence, RI, 2008.

[50] R. Anguelov, C. Dufourd, and Y. Dumont. Mathematical model for pest-insect control using mating disruption and trapping. Applied Mathematical Modelling, 52:437–457, 2017.

[51] R. Anguelov, Y. Dumont, and I.V. Yatat Djeumen. Sustainable vector/pest control using the permanent sterile insect technique. Mathematical Methods in the Applied Sciences, 43(18):10391–10412, 2020.

[52] JS Meza, I ul Haq, MJB Vreysen, K Bourtzis, GA Kyritsis, and C Cáceres. Comparison of classical and transgenic genetic sexing strains of mediterranean fruit fly (diptera: Tephritidae) for application of the sterile insect technique. PLoS ONE, 13(12), 2018.

[53] D.P. Papachristos and N.T. Papadopoulos. Are citrus species favorable hosts for the mediterranean fruit fly? a demographic perspective. Entomologia Experimentalis et Applicata, 132(1):1–12, 2009.

[54] RStudio Team. RStudio: Integrated Development Environment for R. RStudio, PBC., Boston, MA, 2020.

[55] R Core Team. R: A Language and Environment for Statistical Computing. R Foundation for Statistical Computing, Vienna, Austria, 2023.

[56] R.E. Plant and R.T. Cunningham. Analyses of the Dispersal of Sterile Mediterranean Fruit Flies (Diptera: Tephritidae) Released from a Point Source. Environmental Entomology, 20(6):1493–1503, 12 1991.

[57] B.J. Paranhos, N.T. Papadopoulos, D. McInnis, C. Gava, F.S. Lopes, R. Morelli, and A. Malavasi. Field dispersal and survival of sterile medfly males aromatically treated with ginger root oil. Environ. Entomol., 39(2):570–575, 2010.

[58] K. Kraaijeveld, B.I. Katsoyannos, M. Stavrinides, N. A. Kouloussis, and T. Chapman. Remating in wild females of the Mediterranean fruit fly, Ceratitis capitata. Animal Behaviour, 69(4):771–776, 2005.

[59] T. Pogue, K. Malod, and C.W. Weldon. Patterns of Remating Behaviour in Ceratitis (Diptera: Tephritidae) Species of Varying Lifespan. Frontiers in Physiology, 13:824768, 2022.

[60] Diana Pérez-Staples and Solana Abraham. Postcopulatory behavior of tephritid flies. Annual Review of Entomology, 68(1):89–108, 2023.

[61] J. Zhao, S. Chen, Y. Deng, R. He, J. Ma, and F. Liang. Sperm precedence pattern and the effect of irradiation on male mating competition in the oriental fruit fly, bactrocera dorsalis. Biohelikon: Cell Biology, 1:1–5, 2013.

[62] D. Bertin, F. Scolari, C.R. Guglielmino, M. Bonizzoni, A. Bonomi, D. Marchini, L.M. Gomulski, G. Gasperi, A.R. Malacrida, and C. Matessi. Sperm storage and use in polyandrous females of the globally invasive fruitfly, ceratitis capitata. Journal of Insect Physiology, 56(11):1542–1551, 2010.

[63] P.A. Rendón, W.R. Enkerlin, and C. Cáceres. Sterile insect release density calculations spreadsheet. Food and Agriculture Organization of the United Nations/International Atomic Energy Agency. Vienna, Austria. 30 pp., 2019.

[64] A. Bakri, K. Mehta, and D.R. Lance. Sterilizing Insects with Ionizing Radiation. In V A Dyck, Jorge Hendrichs, and A. S. Robinson, editors, Sterile Insect Technique, pages 233–268. Springer, Dordrecht, 2005.

[65] A.M. Costa, C.S. Anjos-Duarte, A.K.P. Roriz, V.S. Dias, and I. S. Joachim-Bravo. Male diet and age influence to inhibit female remating in Ceratitis capitata (Diptera: Tephritidae). Journal of Applied Entomology, 136(6):456–463, 2012.

[66] T.S. Whittier and K.Y. Kaneshiro. Male Mating Success and Female Fitness in the Mediterranean Fruit Fly (Diptera: Tephritidae). Annals of the Entomological Society of America, 84(6):608–611, 1991.

[67] P. Radhakrishnan, D. Pérez-Staples, C.W. Weldon, and P.W. Taylor. Multiple mating and sperm depletion in male Queensland fruit flies: effects on female remating behaviour. Animal Behaviour, pages 1–8, 08 2009.

[68] E. B. Jang. Physiology of mating behavior in Mediterranean fruit fly (Diptera: Tephritidae: Chemoreception and male accessory gland fluids in female post-mating behavior. Florida Entomologist, 85(1):89–93, 2002.

[69] R. Anguelov, Y. Dumont, and J. Lubuma. Mathematical modeling of sterile insect technology for control of anopheles mosquito. Comput. Math. Appl., 64:374–389, 2012.

